# Linking microbial community structure with function using amplicon sequencing of NRPS genes associated with wheat roots under drought stress

**DOI:** 10.1101/2024.08.28.609728

**Authors:** Ying Guan, Edmond Berne, Rosanna Catherine Hennessy, Paolina Garbeva, Mette Haubjerg Nicolaisen, Frederik Bak

## Abstract

Secondary metabolites are bioactive compounds, diverse in structure with versatile ecological functions including key roles in mediating interactions between microorganisms and plants. Importantly, these compounds can promote the colonization of plant surfaces, such as roots, or modulate root exudates to enhance microbial recruitment and establishment. However, owing to the vast diversity of secondary metabolites, their importance in plant root colonization and in particular under stress conditions such as drought, remains unclear. To determine the involvement in root colonization of some of these secondary metabolites, we used amplicon sequencing targeting the adenylation domain of the NRPSs and the 16S rRNA gene from the rhizoplane of wheat grown in soil under normal and drought stress conditions. Results showed that drought transiently affected the bacterial community composition and the NRPS composition in the rhizoplane. We observed that drought selected for distinct groups of siderophores from different taxonomical groups, enriching for *Streptomyces* and depleting *Pseudomonas* siderophores. In addition, drought enriched *Pseudomonas*-derived NRPS genes encoding viscosin, a cyclic lipopeptide with biosurfactant properties, indicating that compounds linked to motility and colonization provide a competitive advantage during rhizoplane colonization under drought stress condition. This observation was experimentally confirmed using the viscosin producing *P. fluorescens* SBW25 and its viscosin-deficient mutant. A higher abundance of SBW25 colonized the roots under drought stress conditions compared to the viscosin-deficient mutant. In summary, our work demonstrates the potential for amplicon sequencing of NRPS genes coupled with *in planta* experiments to elucidate the importance of secondary metabolites in root colonization.

**Importance:** To harness beneficial plant-microbe interactions for improved plant resilience, we need to advance our understanding of key factors required for successful root colonization. Bacterial produced secondary metabolites are important in plant-microbe interactions, and thus, targeting these genes generate new knowledge that is essential for leveraging bacteria for sustainable agriculture. We used amplicon sequencing of the NRPS A domain on the rhizoplane of wheat exposed to drought stress to identify important secondary metabolites in plant-microbe interactions during drought. We show that the siderophores respond differently to drought stress depending on taxonomic affiliation and that the potential to synthesize viscosin increases root colonization. Importantly, this study demonstrates the potential of amplicon sequencing of NRPS genes to reveal specific secondary metabolites involved in root colonization, particularly in relation to drought stress, and highlights how the resolution provided by this approach can link specific compounds to a specific stress condition in a soil system.

## Introduction

Microorganisms are important for plant health and development, as they provide beneficial functions such as nutrient acquisition, disease suppression, or reduction of abiotic and biotic stresses (Mendes *et al*., 2013; Trivedi *et al*., 2020). Secondary metabolites (SMs) produced by rhizobacteria have been proposed to be involved in alleviating plant stress (Chowdhury *et al*., 2015; Lourenzi *et al*., 2022). However, microorganisms produce a broad array of secondary metabolites, and the ecological role of most secondary metabolites in plant-microbe interaction remains largely unknown (Tyc *et al*., 2017).

An important class of secondary metabolites are non-ribosomal peptides (NRPs), synthesized by non-ribosomal peptide synthetases (NRPSs). NRPs have a variety of functions in antimicrobial activity, bacterial motility (de Bruijn *et al*., 2007, 2008), biofilm formation (de Bruijn *et al*., 2008; D’aes *et al*., 2014), root colonization (Tran *et al*., 2007; Guan *et al*., 2024) and the induction of systemic resistance (ISR) in plants (Ma *et al*., 2016; Omoboye *et al*., 2019). These findings are mainly derived from simple systems without soil (but see (Guan *et al*., 2024)), and thus our understanding of the ecological roles of NRPSs in the environment is challenged by a lack of studies targeting the mechanisms of these compounds in complex systems.

We have recently demonstrated that the potential to produce the NRP viscosin enhanced the colonization ability of *P. fluorescens* SBW25 on wheat roots and had a significant impact on shaping the assembly of both rhizosphere and rhizoplane microbial communities (Guan *et al*., 2024). Furthermore, the rhizosphere microbiome contains a distinct composition of NRPSs compared to the neighboring bulk soil (Dror *et al*., 2020) supporting that these compounds play a key role in plant-microbe and microbe-microbe interactions around the roots. While the rhizosphere NRPS of tomato, lettuce, cucumber, potato, and populous have previously been determined (Aleti *et al*., 2017; Blair *et al*., 2018; Dror *et al*., 2020, 2022), these studies primarily focused on finding potential new sources of antimicrobials.

Given the importance of SMs in chemical signalling between microbes and plants, the composition of NRPSs and its temporal dynamics could provide specific cues to mechanisms of NRPs during root colonization. For example, (Aleti *et al*., 2017) showed a change in siderophore composition between emergence and senescence in potato, suggesting that different iron scavenging strategies are advantageous at distinct growth stages. Yet, this is the only study of temporal dynamics of NRPS genes in the root zone.

Secondary metabolites have also been shown to be important in plant-microbe interactions during drought, where beneficial rhizosphere microorganisms are crucial for enhancing plant drought tolerance (Chieb and Gachomo, 2023). Drought stress typically reduces soil microbial diversity, while plants recruit drought-tolerant and plant stress alleviating microbes through complex interactions with their microbiome (Bouskill *et al*., 2013; Preece *et al*., 2019; Igwe *et al*., 2024). Early research showed that microbial exopolysaccharides, associated with biofilm formation, as well as biosurfactants, enhances soil water retention and thus increase plant drought tolerance (Naseem and Bano, 2014; Raddadi *et al*., 2018; Benard *et al*., 2019; Gutierrez *et al*., 2022). However, no studies have addressed the effect of drought stress on the development in composition of the NRPSs at the root surface. Such studies are needed to disentangle the impact of NRPs in drought resistance.

NRPs are synthesized by modular enzymes called NRPSs. Each module incorporates a specific amino acid into the peptide chain, starting with the adenylation (A) domain which selects and activates the amino acid. The specificity of the A domain is a key determinant in peptide sequence and function and can be targeted through amplicon sequencing. Recently, culture-independent approaches, and in particular amplicon sequencing of NRPS gene clusters, have shed light on how these compounds shape the microbial communities in the rhizosphere (Aleti *et al*., 2017; Tracanna *et al*., 2021). Amplicon sequencing has not been limited to NRPS, but includes polyketide synthases, and have been used to describe the composition of these groups in lettuce and tomato (Dror *et al*., 2020), and in different environments (Geers *et al*., 2023).

Here, we studied the rhizoplane bacterial community and NRPS composition during the early growth stages of wheat under drought stress and subsequent recovery. Using amplicon sequencing of the 16S rRNA gene and the NRPS A domain, we specifically aimed to 1) characterize the NRPS diversity and composition in the rhizoplane of well-watered wheat plant, 2) analyze the temporal dynamics of NRPS composition in the rhizoplane and 3) assess the effect of drought on the NRPS composition in the rhizoplane.

## Results

### Drought transiently impacts bacterial community taxonomic composition and diversity

We grew winter wheat (cv. Sheriff) for a duration of 5 weeks under controlled conditions in a growth chamber. The soil was collected from a nearby agricultural field and mixed with sterile sand (3:1; from here on “soil”). Drought was imposed on 2-weeks old wheat seedlings, by not watering the plants for two weeks, where after water was added to alleviate the drought stress (Fig S1). Control plants received continued watering throughout the entire experiment.

The drought-stressed plants showed reduced shoot length, shoot fresh weight and root dry weight 4 weeks post sowing (Fig 1A-C). In addition, a reduction in chlorophyll content (t-test, p < 0.001), as well as higher peroxidase dismutase (POX) (t-test, p < 0.001) was observed (Fig. S2). No difference in superoxide dismutase (SOD) activities was recorded as compared to control plants (Fig S2). At 5 weeks, following one week of re-watering, drought-stressed plants remained growth impaired, but showed signs of recovery in shoot fresh weight (Fig 1A-C). Taken together, this shows that the treatment did impose a drought stress response during the experiment.

**Figure 1.**
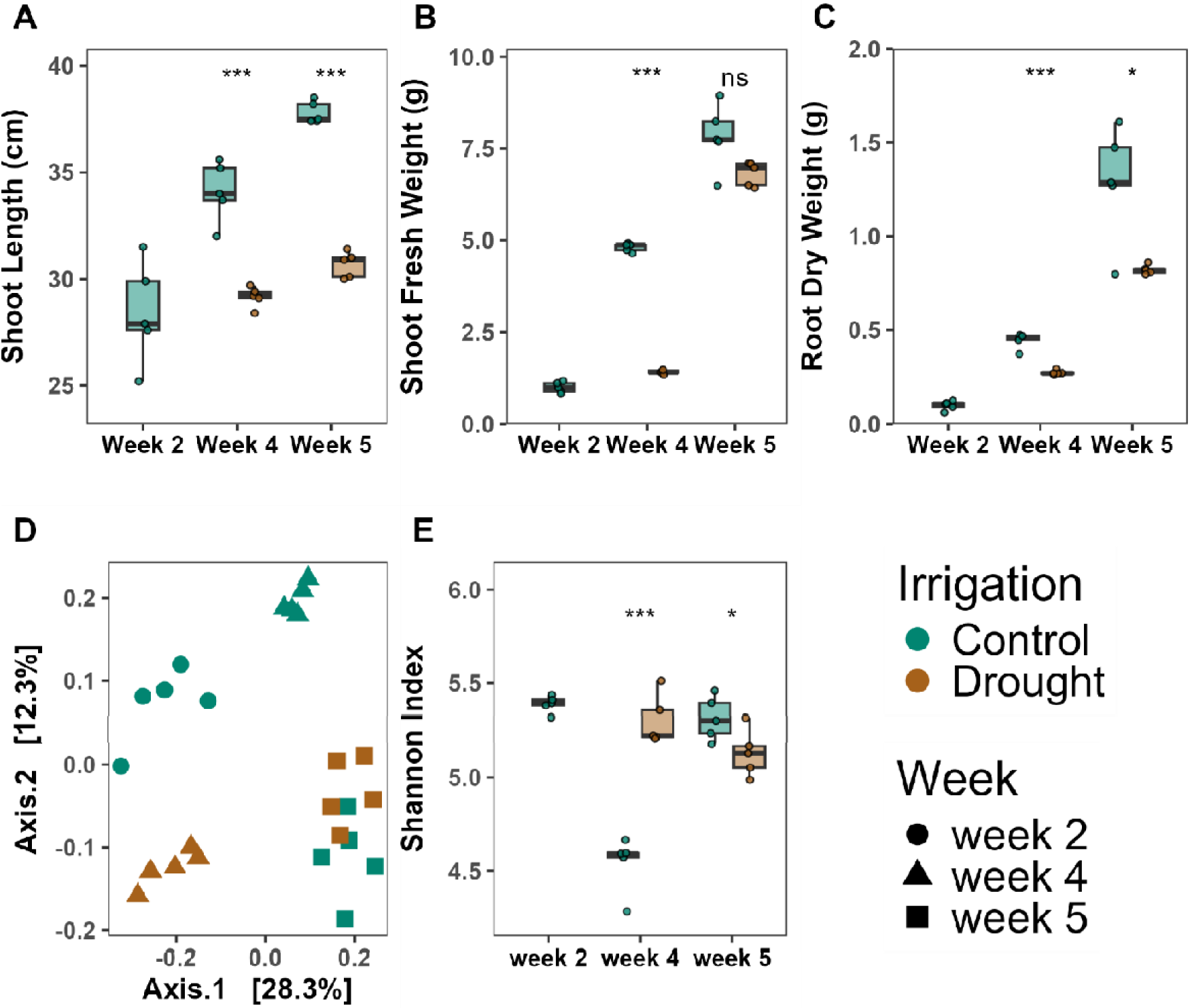
The effect of drought on plant growth before, during, and after drought. The shoot length (A), shoot fresh weight (B) and root dry weight (C) at different sampling times after sowing. Control: well-watered plants, Drought: drought stress plants. (D) PCoA of Bray Curtis dissimilarities of the bacterial community based on 16S rRNA amplicon sequencing. (E) Shannon diversity. Each box plot represents data from five replicates. Each dot represents a sample point. Asterisks in A-D indicate a statistically significant difference between the two groups (Drought and Control), which was derived from the t-test: \**p* < 0.05, \*\**p* < 0.01 and \*\*\**p* < 0.001. The horizontal bars within boxes represent medians. The tops and bottoms of boxes represent the 75th and 25th percentiles, respectively. The upper and lower whiskers extend to data no more than 1.5× the interquartile range from the upper edge and lower edge of the box, respectively.

To determine the effect of the drought stress on bacterial rhizoplane communities, we initially performed 16S rRNA gene amplicon sequencing targeting the V5-V7 regions. We obtained 1,691 Amplicon Sequence Variants (ASVs) with a median sequencing depth of 19,328 reads per sample, with a range of 12,905 to 28,215 reads per sample. Rarefaction curves indicated that sufficient sequencing depth was reached (Fig. S3). Drought changed the community composition (Fig 1D, Table S1), but this change was transient as communities after 5 weeks i.e., 1 week of re-watering, clustered together, as indicated by the positive interaction between time and water level (PERMANOVA, R^2^ = 0.11, p = 0.001, Table S1).

Concomitantly, drought resulted in higher Shannon diversity in the rhizoplane community compared to control plants (p < 0.001, t-test) after 4 weeks, whereas the Shannon diversity was higher in the control plants at week 5 (p = 0.043, t-test) (Fig. 1E).

At the ASV level, 48 ASVs increased in relative abundance during drought, while nine ASVs were more abundant in control plants (Corncob, FDR < 0.05, Fig. S4). Several bacterial genera, including *Bacillus, Streptomyces* and *Kribella* increased in relative abundance on the rhizoplane in the drought-stressed plants. In contrast, only three genera, *Massilia, Flavobacterium,* and *Arenimonas,* decreased in relative abundance during drought (Fig. S5). At week 5, only 7 genera were differential abundant between the two treatments (Fig. S6).

In conclusion, drought-induced plant effects were sustained throughout the experiment, while effects on the associated bacterial communities were transient, with a converging of bacterial communities for both treatments at week 5.

### Temporal dynamics is a key driver of NRPS composition and diversity

To go beyond taxonomic characterization of the microbiome, we performed amplicon sequencing targeting the NRPS A domain (Tracanna *et al*., 2021) from the rhizoplane at the three sampling points for both drought stressed and non-drought (control) plants. After processing of reads, NRPS sequences were clustered into Amplicon Clusters (ACs) as in (Tracanna *et al*., 2021). The 25 samples contained 2,674,962 reads, and 33,015 ACs. The median number of reads was 105,643 per sample, with a range of 22,354-230,381 reads. Rarefaction curves showed that the majority of the diversity was captured at around 40,000 reads per sample (Fig. S7). Neither the alpha diversity (Shannon index) (Fig. 2A) nor the richness of the NRPS domain A amplicons differed between the sampled time points in the control plants. In contrast, developmental stage was a main driver of the NRPS domains composition (PERMANOVA, R^2^ = 0.23, p =0.001, Fig. 2B, Table S2) in the control plants, explaining 23% of the variation.

**Figure 2.**
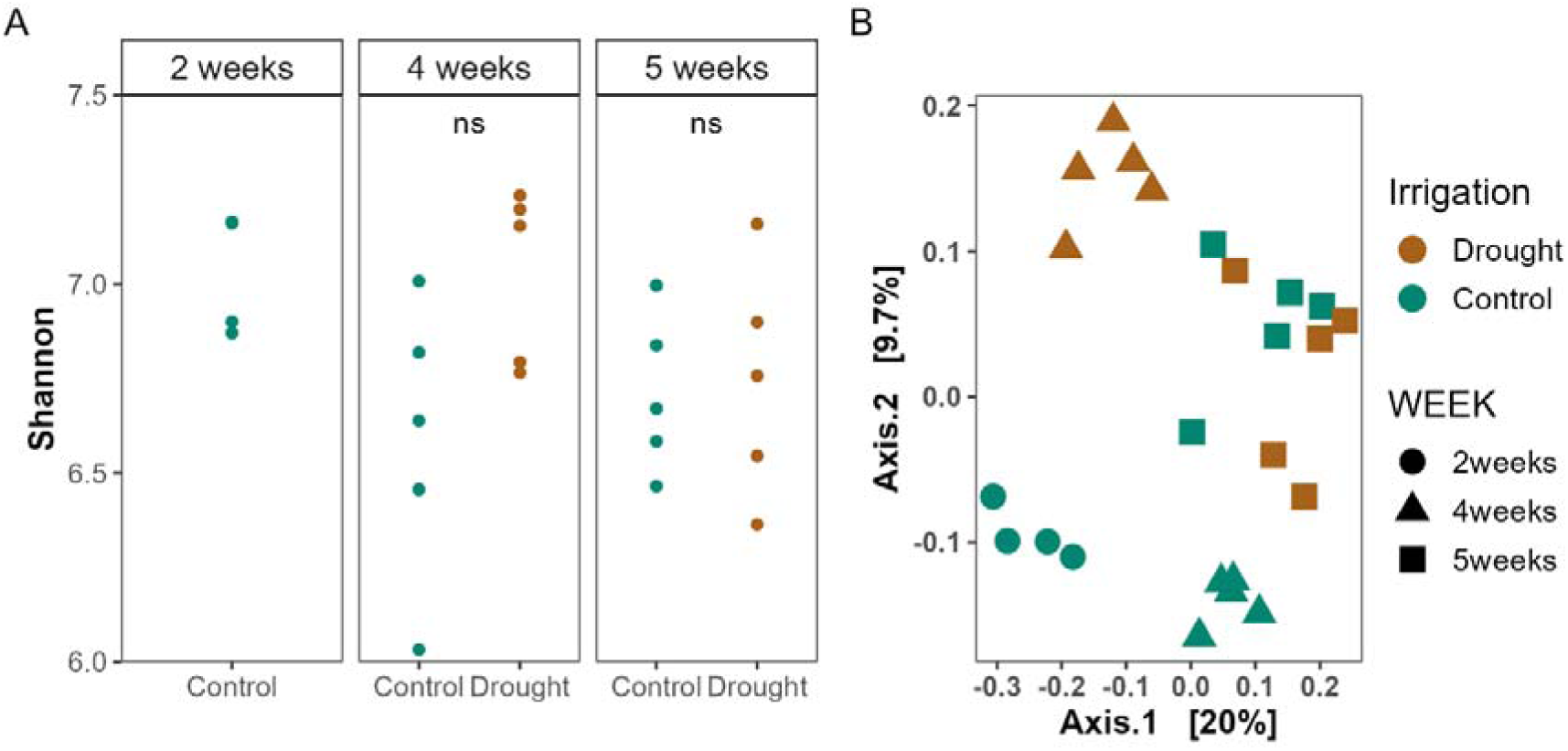
A) Shannon index and B) PCoA of rarefied NRPS ACs from the rhizoplane based on Bray-Curtis dissimilarities. NS = not significant (p > 0.05, t-test).

### Drought alters NRPS composition to a lower extent than bacterial composition

As the 2-week drought period changed the bacterial community in the rhizoplane, we investigated whether this observation was also applicable for the NRPS composition. Drought did not change the Shannon diversity (Fig. 2A) or the richness (Fig. S8) either at week 4 or week 5, contrasting the findings from the 16S rRNA amplicons. In line with observations at the taxonomic level, drought changed the composition of the NRPS genes (Fig. 2B) (PERMANOVA, R^2^ = 0.08, p = 0.005). Further, the interaction between developmental stage and drought was also significant (PERMANOVA, R^2^ = 0.09, p = 0.001), indicating that the effect of drought on the NRPS domain composition was transient. Based on visual inspection of the PCoA, the drought and control samples clustered together at week 5 supporting that the effect of drought was transient (Fig. 2B). A comparison of the NRPS dissimilarity matrix with the 16S rRNA dissimilarity matrix showed an association between the two (Mantel’s r = 0.453, p = 0.001). Hence, only 45% of the variation in NRPS domain A diversity can be explained by the 16S rRNA taxonomic marker.

### Actinomycetota and Proteobacteria represent the largest proportion of NRPS ACs

Next, the NRPS reads were annotated using MiBig and Antismash databases in the dom2bgc pipeline (Tracanna *et al*., 2021) with the aim of assigning both taxonomy and function to the ACs. Despite using the most recent databases, only 15% and 16% of the ACs were annotated as NRPSs in the control plants and the drought treated plants, respectively, across the three sampling times. Due to the high percentage of unassigned reads, we mapped the reads back to the model strains *P. fluorescens* SBW25. All reads mapping to the SBW25 genome aligned with known NRPS biosynthetic gene clusters, giving high fidelity in the primers used.

Reads were taxonomically assigned to seven different phyla, with the majority (64-73%) of the ACs assigned to Actinomycetota, with Proteobacteria and Myxococcota comprising 18-24% and 8-13% depending on the watering regime, respectively (Fig. 3A). Together, Bacillota, Gemmatimonadota and Verrucomicrobiota comprised ∼1% of the annotated reads. For Bacillota, reads were assigned to *Paenibacillus, Thermoactinomyces, Tumebacillus* and *Anoxybacillus*, whereas reads were only assigned to *Longimicrobium* and *Luteolibacter* for Gemmatimonadota and Verrucomicrobiota, respectively. NRPS genes annotated to Planctomycetota (genus: *Aquisphaera*) were only detected in the drought-stressed plants at week 4 (<0.05% of annotated ACs).

**Figure 3.**
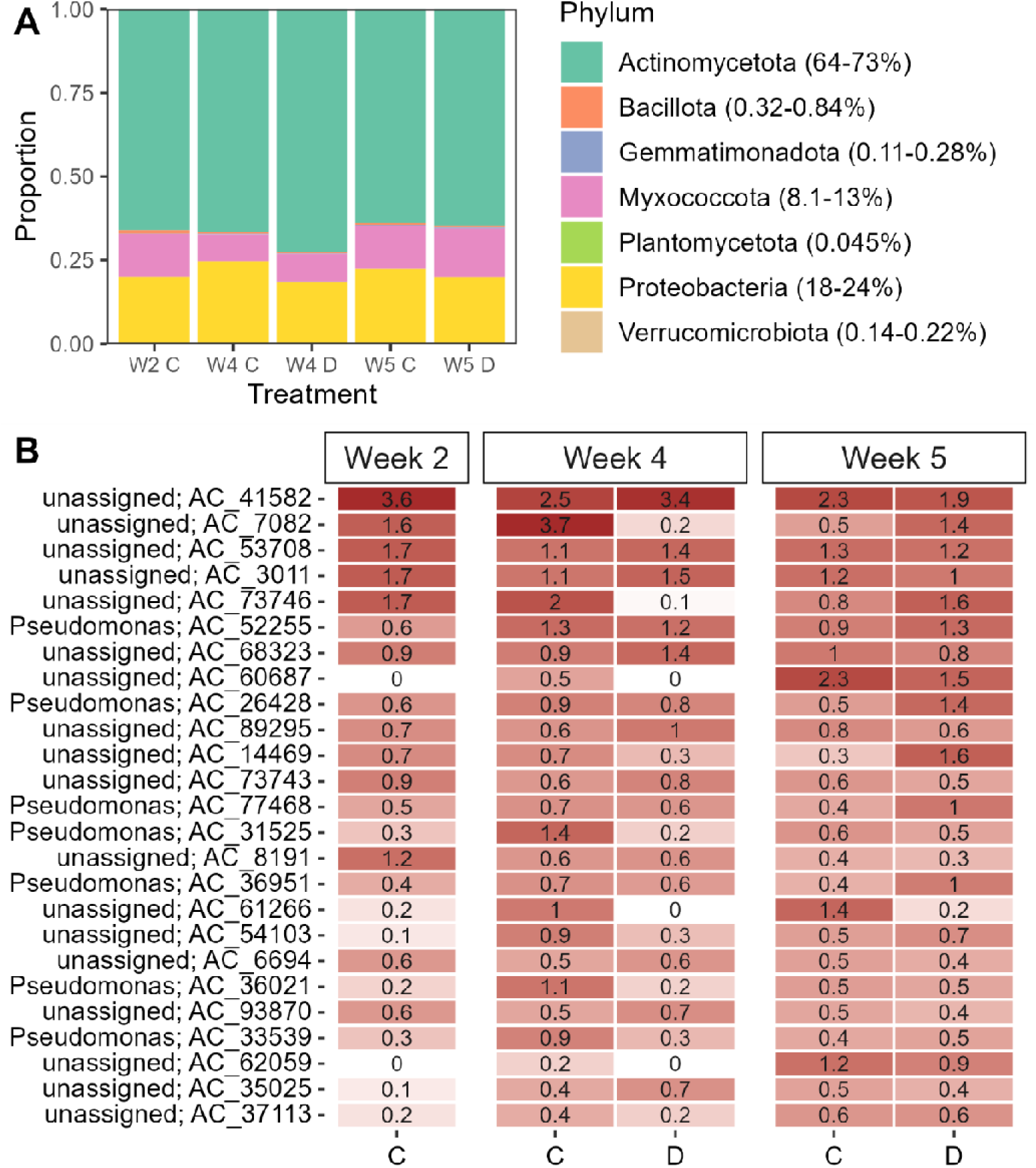
A) Taxonomic assignment at the phylum level of the ACs in drought stressed plants and control plants. Numbers in parentheses indicate range of the relative abundance across samples. B) Mean relative abundance of the 25 most abundant ACs across all sampling times and treatments in the rhizoplane (n = 5). C = Control, D = Drought.

Of the 25 most abundant NRPS ACs in the rhizoplane across developmental stage, seven were taxonomically assigned to *Pseudomonas* (Fig. 3B), and functionally classified as NRPS. The rest were unassigned. Due to the low proportion of assigned amplicon clusters, we used blastp to search the 25 most abundant ACs in the MiBig database v.3.1 (Table S3). The most abundant (amplicon_cluster_41582) was classified as Pf-5 pyoverdine. Of the *Pseudomonas* associated NRPS reads, six of the seven were assigned to pyoverdine-related “compounds” (Table S3). Despite our effort, 10 of the 25 most abundant amplicon clusters did not represent known biosynthetic gene clusters indicating a potentially high and untapped abundance of novel compounds in the rhizoplane.

To determine which ACs were enriched or depleted on a temporal scale, we tested differentially abundant ACs in the control plants using corncob (Martin *et al*., 2020). This showed that histicorrugatin, pyoverdines and unassigned ACs from *Pseudomonas* increased from week 2 to week 4 (Fig. S9). Pyoverdine, not assigned to any taxa, together with bacillibactin and occidiofungin decreased between week 2 and week 4 (Fig S9). In total, 108 ACs changed in relative abundance from week 2 to week 4. In contrast, only 39 ACs differed in relative abundance between week 4 and week 5 indicating a less dynamic functional microbiome at later developmental stages. Histicocorrugatin and pyoverdine from *Pseudomonas* together with an unassigned phenalamide decreased from week 4 to 5 (Fig. S10), while an omnipeptin assigned to *Saccharotix* increased in relative abundance. However, the majority of ACs that increased were either taxonomically or functionally unassigned.

### Drought reduces the relative abundance of *Pseudomonas* pyoverdine related NRPS genes in the rhizoplane

We then asked which ACs were enriched or depleted when plants were exposed to drought. In week 4, drought stress increased the relative abundance of 64 ACs and decreased the relative abundance of 37 ACs (Fig. S11). The majority of ACs were unassigned using the dom2bgc pipeline among the differentially abundant ACs at week 4. However, we identified eight ACs taxonomically assigned to *Streptomyces*, one assigned to *Rhodococcus*, one assigned to *Lentzea,* and two assigned to *Saccharotrix* among the ACs enriched in the drought stressed plants at week 4. Contrastingly, we found seven *Pseudomonas* assigned ACs, one *Mycolibacterium* assigned AC and two *Pseudoxanthamonas* assigned ACs that were depleted during drought (Fig. S11). They were functionally annotated with a blastp search against the MiBig database, and we identified genes encoding for bacillibactin, peuchelin and potashchelin as ACs with highest prevalence among the differentially abundant ACs in the drought-stressed rhizoplane samples (Fig. 4, Table S4). These compounds are siderophores taxonomically assigned to *Bacillus, Streptomyces* and *Halomonas*. Conversely, *Pseudomonas* genes coding for siderophores, such as pyoverdine, and histicocorrugatin decreased in relative abundance during drought (Fig. 4, Table S4). One week later, 11 AC were enriched and 11 AC were depleted in the drought treated plants as compared to the control plants (Fig. S12). Most of the ACs with a relatively higher abundance in the drought-stressed plants in week 5 were genes encoding for pyoverdines.

**Figure 4.**
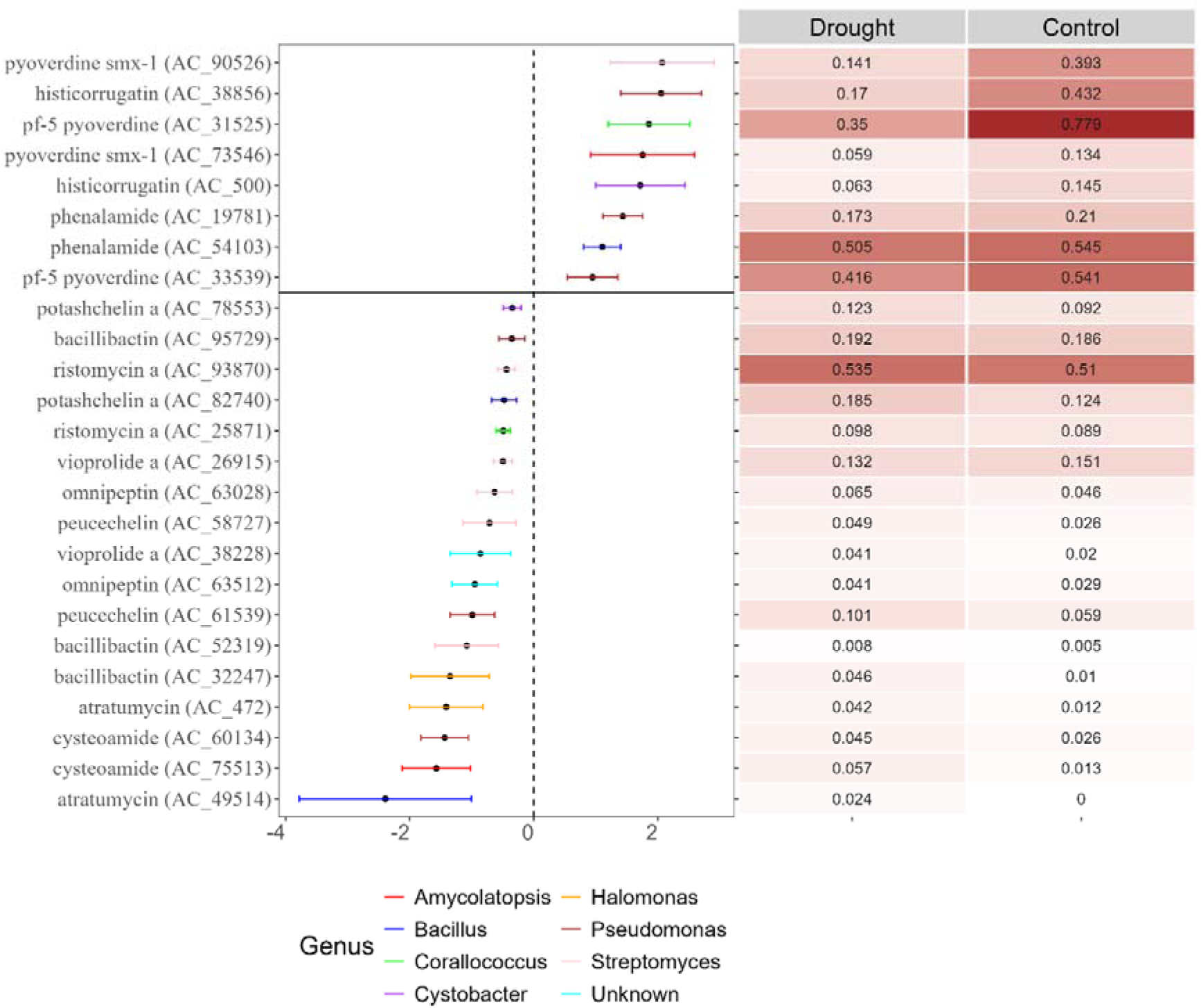
Differential abundance test (corncob, FDR < 0.05), right side shows the ACs depleted during drought, and left side shows the ACs enriched during drought. Mean relative abundances of the ACs in the rhizoplane at week 4 are shown in the heatmap on the right (n=5) Only ACs that were annotated using MiBig and found at least twice are shown. See Fig. S11 for a full list of differential abundant ACs.

To complement this analysis, we identified core ACs that were only present in drought-stressed or control plants (unique ACs). Here, we identified that the drought-stressed plants hosted a higher number of unique ACs (10) compared to the control/watered plants (3 unique ACs) at the end of the drought period (week 4) (Table 1). Cooperating the results from the differential abundance test, coechelin encoding NRPS genes was also among the unique ACs in the drought-stressed plants at week 4. Coechelin is a siderophore encoded by *Streptomyces*. Together, this shows that siderophores produced by *Streptomyces, Bacillus* and *Halomonas* are enriched during drought while *Pseudomonas* produced siderophores are depleted. This suggests that different iron scavenging strategies are decisive for success in root colonization during drought.

**Table 1.** MiBig annotation of core ACs unique to either the drought stressed or control plants at week 4. There was also an unknown AC among the unique ACs in the drought stressed plants.

| Drought | Compound Class | Organism | Mode of Action | References |
| --- | --- | --- | --- | --- |
| Scytocyclamide | cyclic lipopeptide | *Scytonema hofmannii* | Antimicrobial activity | (Heinilä *et al.*, 2020) |
| Coelichelin | peptide siderophore | *Streptomyces coelicolor* | Iron Scavenging | (Challis and Ravel, 2000) |
| Aurantimycin | cyclic lipopeptide | *Streptomyces aurantiacus* | Antimicrobial activity | (Schlegel *et al.*, 1995; Zhao *et al.*, 2016) |
| Polyoxypeptin | cyclic depsipeptide | *Streptomyces* sp. | Activity against cancer cells | (Umezawa *et al.*, 1998) |
| Ohmyungsamycin | cyclic depsipeptide | *Streptomyces* sp. | Antimicrobial activity | (Hur *et al.*, 2018) |
| Pepticinamin | cyclic lipopeptide | *Actinobacteria bacterium* | Antimicrobial activity | (Santa Maria *et al.*, 2019) |
| Myxochromide | cyclic peptide | *Pseudomonas putida* | Antimicrobial activity | (Stephan *et al.*, 2006) |
| Althiomycin | pentapeptide | *Myxococcus xanthus*, *Serratia marcescens* | Antimicrobial activity | (Cortina *et al.*, 2011; Gerc *et al.*, 2012) |
| Rotihibin | cyclic lipopeptide | *Streptomyces* sp. | Herbicidal | (Halder *et al.*, 2018; Planckaert *et al.*, 2021) |
| Control |  |  |  |  |
| Myxothizaol | respiratory chain inhibitor | *Myxococcus fulvus* | Antifungal | (Thierbach and Reichenbach, 1981) |
| Thiocoraline | bicyclic peptide | *Verrucosipora* sp. | Cytotoxic against cancer cells | (Wyche *et al.*, 2011) |
| Necroxime | cyclic lipodepsipeptide | *Mortierella verticillata* | Antinematocidal | (Büttner *et al.*, 2021, 2022) |

### The NRPS-derived CLP viscosin produced by *Pseudomonas* enhances root colonization under drought stress

NRPs previously reported to play key roles in root colonization include CLPs produced by *Bacillus* and *Pseudomonas* (Bais *et al*., 2004; Tran *et al*., 2007; Aleti *et al*., 2016; Guan *et al*., 2024). However, this has only been demonstrated during well-watered conditions. To determine whether CLPs provide an advantage in root colonization under drought stress we mined the NRPS amplicons for CLPs produced by *Pseudomonas* (Zhou *et al*., 2024).

*Pseudomonas* comprised the highest proportion of the identified CLPs with a taxonomic assignment in the rhizoplane at all sampling times and water treatments (Fig. S13). The control plants harboured 2.5 times higher proportion of *Pseudomonas* CLP-related ACs than drought stressed plants at 4 weeks, while this difference was reduced to 0.5 times higher at week 5. This suggests that on a general note, CLPs are not selected for under drought stress. As we previously showed that the potential to synthesize viscosin provides an advantage in root colonization for the *P. fluorescens* SBW25 (Guan *et al*., 2024), we were particularly interested in the NRPS ACs linked to this compound. Viscosin, together with viscosinamide, was one of the most relative abundant *Pseudomonas* CLPs found across all samples in the rhizoplane (Fig. 5) and the most abundant at week 2. While viscosin coding ACs proportion of all ACs were comparable between treatments at week 4, viscosin coding ACs comprised 10% (0.048/0.47) of the *Pseudomonas* CLP-encoding ACs in the control plants and 22% (0.044/0.20) in the drought stressed plants. This difference was even more noticeable after 5 weeks, where the viscosin coding ACs comprised 26% (0.045/0.18) of the *Pseudomonas* CLP encoding ACs in the drought-stressed plants compared to 5.6% (0.15/0.27) of these CLPs in the control plants. The increased proportion of viscosin-related ACs in drought stressed plants indicates that *Pseudomonas* harbouring the viscosin biosynthetic gene cluster colonize the rhizoplane better than other CLP producing pseudomonads. Thus, we hypothesized that the potential to produce viscosin would increase a strains’ root colonization potential during drought.

**Figure 5.**
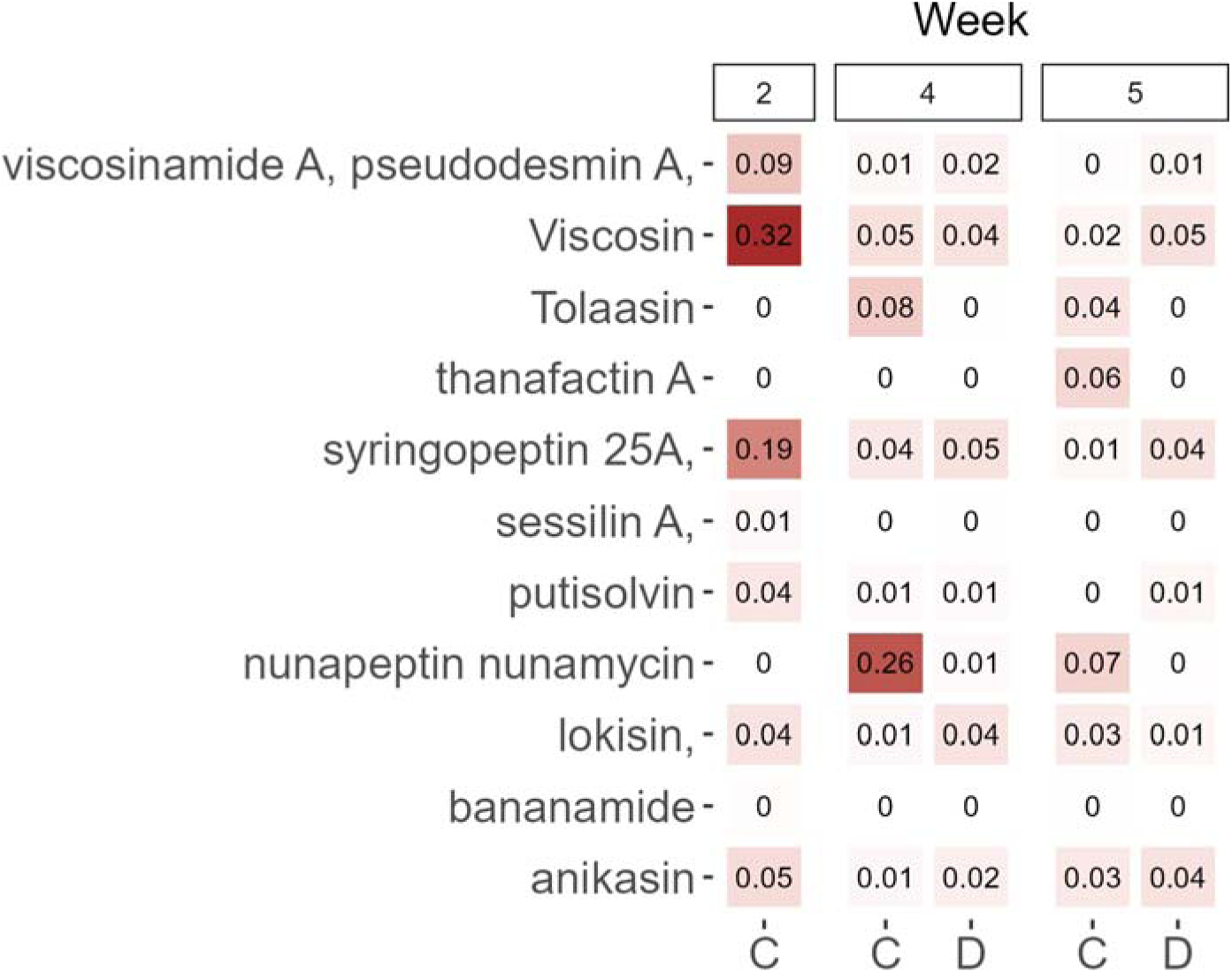
Proportion of reads assigned to *Pseudomonas* CLPs out of the total number of reads in the NRPS amplicon data. Values are mean (n=5). C = Control, D = Drought. Numbers above indicate week of sampling.

To test this hypothesis, we inoculated surface-sterilised seedlings with a viscosin producing strain *P. fluorescens* SBW25 (WT) and a *viscA* deficient mutant (Δ*viscA*) unable to synthesize viscosin, and grew the plants as described above. We have previously shown that the WT colonizes wheat roots better under well-watered conditions (i.e. control plants) using both qPCR, CFU counts and microscopy (Guan *et al*., 2024). Both WT and Δ*viscA* were chromosomally tagged with mCherry gene, which we targeted with qPCR. In both control and drought stressed plants, we observed a higher abundance of mCherry genes per gram of rhizoplane soil in the WT inoculated plants compared with plants inoculated with the Δ*viscA* mutant after 4 weeks (Fig. 6). The number of mCherry genes decreased from week 4 to week 5, but the WT inoculated plants still showed higher colonization compared to the Δ*viscA* in drought stressed plants. At both sampling points, the mCherry abundances suggest a higher colonization of the WT in the drought stressed plants compared to the control plants. Whether the increased abundance of SBW25 compared to Δ*viscA* at week 4 had an impact on plant health was determined using chlorophyll, SOD and POX activity measurements (Fig. S2).

**Figure 6.**
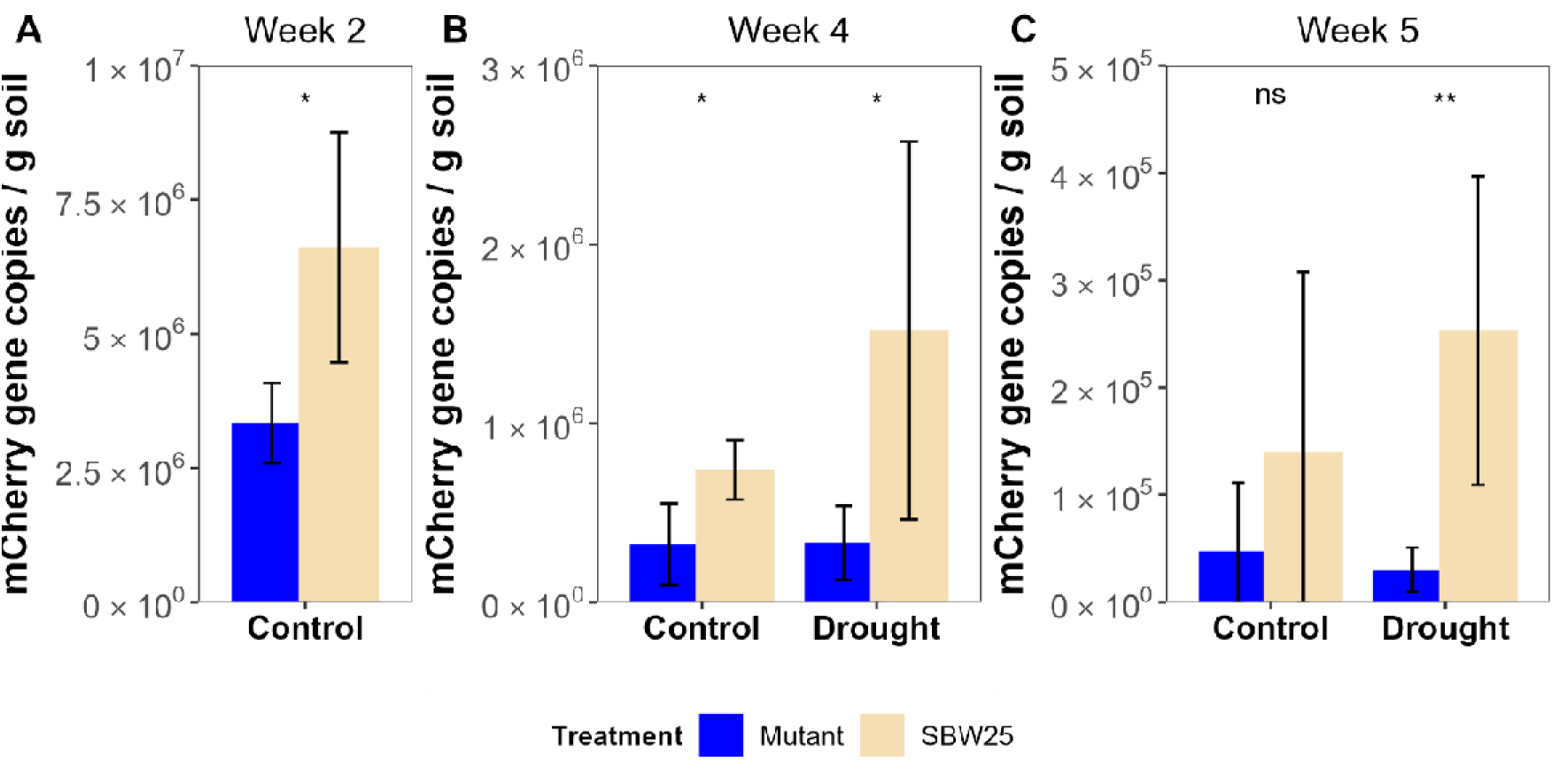
Absolute abundances of mCherry gene copies in the rhizoplane, at week 2 (A), week 4 (B) and week 5(C). The height of the bars represents the mean value. Error bars are std. dev based on five biological replicates. Asterisks indicate a statistically significant difference between the two inoculations (t-test *p < 0.05 and **p < 0.01).

For control plants, there were no differences among inoculations for chlorophyll or SOD and POX activities. In drought stressed plants, WT inoculation increased the chlorophyll content compared to Δ*viscA* inoculated (p = 0.04, t-test) and non-inoculated plants (p < 0.001, t-test), and was close to the content in plants grown under controlled conditions. In addition, WT inoculation increased the SOD activity compared to non-inoculated plants (p = 0.037, t-test), but not the Δ*viscA.* Inoculation with WT or Δ*viscA* did not impact the POX activities. Hence, the inoculation with WT seemed to alleviate plant stress symptoms of low chlorophyll content. At the microbial side, this experiment supports the hypothesis that the ability to produce viscosin provides a competitive advantage in root colonization not only under normal conditions in natural soils, highlighting its general importance for plant root colonization by *Pseudomonas* species. Furthermore, it supports a role of viscosin during post-drought recovery. Additionally, we conclude that not all CLPs results in an improved colonization during drought.

## Discussion

Bacterial produced NRPs play important roles in plant-microbe interactions (Jacoby *et al*., 2021). To date, research NRPs in relation to plant growth have primarily focused on their growth promoting activities or their biocontrol activity (Zhou *et al*., 2024). However, their role in root colonization and specifically during drought stress have received limited attention.

Here, we exposed wheat (cv. Sheriff) to drought and determined the taxonomic and functional composition in the rhizoplane before, during and after the drought period. We found that drought stressed plants depleted NRPS genes encoding for *Pseudomonas* siderophores, while genes encoding for *Streptomyces* siderophores increased in relative abundance in drought stressed plants. In contrast, drought increased the relative abundance of biosynthetic genes coding for viscosin, a CLP produced by pseudomonads. Secondly, we therefore hypothesized that viscosin confers an advantage in root colonization under drought stress. We experimentally validated this using a natural soil system to compare colonization under drought stress of a viscosin producing wild-type model strain *P. fluorescens* SBW25 with its viscosin-deficient mutant.

Overall, we observed that the NRPS composition in the rhizoplane changed over time. This has also been shown in the rhizosphere of potato (Aleti *et al*., 2017) and reflects the change in overall community composition, as determined by 16S rRNA genes, and presumed changes in root exudates which attracts bacteria. Nonetheless, the change in NRPS composition also indicates that the genes required for root compatibility changes over time. Especially, NRPs such as pyoverdines and histicorrugatin (*Pseudomonas* siderophores) changed with plant developmental stage, peaking at week 4, indicating that the form of available iron around the roots changes with growth stage. This corroborates the findings by Aleti *et al* (2017), who also identified the relative abundances of siderophores to be dynamic at two different growth stages in potato.

A 2-week drought period changed the bacterial community, both at the taxonomic (16S rRNA gene) and the functional level (NRPS A-domain). However, drought did not affect the alpha diversity and the richness of the NRPSs. Most of the NRPS ACs were unannotated, as also demonstrated in other studies applying NRPS or PKs amplicon sequencing in environments such as soil and rhizosphere (Dror *et al*., 2020; Tracanna *et al*., 2021; Geers *et al*., 2023). Due to the large proportion of unassigned ACs in the dataset using the dom2bgc pipeline, we relaxed the cut-off criteria using blastp against the Mibig database. Through this we identified two groups of NRPS related gene groups that were affected by drought.

One group of NRPS genes affected by drought encoded iron-scavenging siderophores. *Pseudomonas-*encoded pyoverdines decreased during drought, whereas *Streptomyces* encoded coechelin, but also siderophores encoded by *Halomonas* and *Bacillus* increased as a result of the drought. The latter coincided with an increase in relative abundance of *Streptomyces* in the 16S rRNA gene amplicons. A longer drought period increases the aerobic conditions in the soil, which in turn oxidises the iron to form insoluble iron oxides (Bouskill *et al*., 2013), reducing the availability for plants and microorganisms. The decrease of *Pseudomonas* genes encoding for siderophores then suggests that they are not able to dissolve these insoluble iron oxides, in contrast to *Bacillus, Halomonas* and *Streptomyces*. In addition, *Pseudomonas* can scavenge iron from phytosiderophores (Jurkevitch *et al*., 1993), which are reduced by plants during drought. This has led Xu et al. (2021) to propose that the reduction of plant phytosiderophores during drought favours *Streptomyces* (or Actinobacteria in general) around the roots of sorghum (Xu *et al*., 2021). This might be at play around the roots of wheat as well during drought, indicating that this is a common phenomenon in plant-microbe interactions. In support of this, when plants were rewatered in week 5, the relative abundance of several ACs coding for *Pseudomonas*-encoded pyoverdines increased. While the work of Xu *et al*. (2021) was performed using shotgun metagenomics, we here demonstrate the potential of using NRPS amplicons for inferences of the ecology in the root zone. Furthermore, this suggests that a relaxed cut-off criteria during blastp was reasonable.

The second group of NRPS genes affected by drought were CLPs, a group of metabolites important in plant-microbe interactions (Zhou *et al*., 2024). Species within *Pseudomonas* produce a vast collection of these compounds, as they have a role in biofilm dispersal, as antimicrobials and in root colonization (Zhou *et al*., 2024). Here, our analyses of the NRPS indicated that the ability to produce the CLP viscosin is an important trait for root colonization of wheat by *Pseudomonas* during drought. This was supported by the higher abundance of *P. fluorescens* SBW25 in the rhizoplane during and after exposure to drought compared to its viscosin deficient mutant. In this work, we determined the abundance using qPCR targeting the mCherry gene, as this detection method was supported by CFU counts and microscopy images in a comparable setup (Guan *et al*., 2024). Viscosin likely functions as a wetting agent capable of enhancing solubility of nutrients and hydrophobic substrates, as proposed for other CLPs (Nybroe and Sørensen, 2004). For example, surface-active *P. fluorescens* and *P. putida* strains can increase leaf wetness and may enhance leaf attachment (Bunster *et al*., 1989). Furthermore, syringafactins from the plant pathogen *P. syringae* can attract water vapor, thereby alleviating water stress on dry leaves (Burch *et al*., 2014). Thus, we speculate that *P. fluorescens* SBW25 uses viscosin as a wetting agent to enhance root attachment and increase nutrient availability. Whether this nutrient availability benefits the plant needs to be determined, but the inoculation with SBW25 increased chlorophyll content and the POX activity of the leaves after 4 weeks during drought stress. An increase in POX after inoculation with PGBR during drought stress has been seen before in lettuce (Kohler *et al*., 2008) and jujube (Zhang *et al*., 2020).

On a broader level, the composition of 16S rRNA gene and NRPS amplicons converged one week after stopping the drought period. Rewetting after drought increases microbial biomass in the rhizosphere (Karlowsky *et al*., 2018), as a result of accumulated dissolved organic carbon during drought (Canarini *et al*., 2017). Further, (Karlowsky *et al*., 2018) found that both gram positive and gram negative bacteria took up ^13^C labelled carbon to the same extent after rewetting. These studies support our findings that rhizosphere bacteria respond evenly in carbon uptake to rewetting. However, at a finer resolution, we still saw a “legacy” effect of drought on the viscosin encoding ACs, which were relatively more abundant one week after stopping the drought, as well as on other pseudomonads CLPs. This points to an underlying functional microdiversity, e.g. the ability to produce viscosin, in the microbial community supporting different ecotypes within the same species (García-García *et al*., 2019). This is supported by the fact that only 45% of the taxonomic diversity explains the NRPS diversity observed. In summary, the effects of drought on the community level were transient, but the NRPS amplicons revealed that functionality was affected for a longer duration. Hence, new knowledge on the microdiversity beyond the taxonomic level is needed to gain a complete understanding of microbial interactions in the plant-soil interface.

Our data analysis identified numerous unassigned ACs highlighting the need for further research in this area to strengthen currently available databases. Interestingly, NRPSs assigned to Verrucomicrobiota, Gemmatimonadetes and Planctomyceota were recently linked to the synthesis of secondary metabolites in soil (Crits-Christoph *et al*., 2018; Sharrar *et al*., 2020), suggesting that they contribute an important diversity of secondary metabolites in the rhizosphere. Despite this inherent limitation, our study demonstrates the potential of amplicon sequencing of NRPS genes to reveal specific secondary metabolites involved in root colonization and in relation to drought stress which can be supported with experimental work.

Overall, our study shows that the root microbiome harbours a wide range of NRPS-genes changing in composition during plant growth and under drought stress. Specifically, the differences in siderophore encoding genes highlights the importance of identifying ecosystem-associated keystone metabolites. Furthermore, we demonstrated how an NRPS amplicon approach can be used to identify key metabolites, e.g. viscosin, in *Pseudomonas* required for root colonization during drought, and further confirmed this finding by experimental work. Expanding our current repositories of NRPS clusters will enhance the use of these amplicons to resolve important ecological questions on the role of NRPs in shaping root colonization patterns and mediating plant-microbe interactions in the rhizosphere.

## Materials and Methods

### Plant materials and growth conditions

The winter wheat cultivar Sheriff (Sejet Plant Breeding, Horsens, Denmark) seeds were used in this study. Initially, the seeds were submerged in sterile Milli-Q water for an hour and subsequently relocated to Petri dishes with dual layers of sterile filter paper, saturated with 5 ml of sterile Milli-Q water for germination. For optimal germination, the seeds were kept in the dark at room temperature for three days. Following this, the three-day-old germinated seedlings were root-dipped in sterile saline solution (0.9% NaCl) and transplanted into polyvinyl chloride (PVC) pots (24 cm in height and 7 cm in diameter) with drainage holes, with each pot accommodating one plant. The pots were filled with approximately 1000g of non-sterile field soil premixed with sterile sand (DANSAND, Brædstrup, Denmark) at a 3:2 ratio (“soil” from here on). The field soil was gathered from the plough layer (0 to 25 cm) at the University of Copenhagen’s experimental farm in Taastrup, Denmark (55° 40′N, 12° 17′E).

The water content of the soil was maintained at 17% (w/w). Following the transplantation of the germinated seeds into the pots, the soil was supplemented with a nutrient solution Nitrogen – Phosphorous – Potassium (NPK 3-1-4) (Drivhusgødning, Park^®^). Each pot received 0.83 ml of the nutrient solution, comprising 2.04% nitrate nitrogen (N), 0.3% ammonium nitrogen (N), 0.46% amide nitrogen (N), 0.69% phosphorus (P), 4.38% potassium (K), 0.08% sulfur (S), 0.06% magnesium (Mg), 0.033% iron (Fe), 0.013% manganese (Mn), 0.002% copper (Cu), 0.002% zinc (Zn), 0.0006% molybdenum (Mo), and 0.004% boron (B). Subsequently, the pots were positioned in the growth chamber that was set at 600 μmol m^−2^ s^−1^ light intensity, 60% humidity, a 16/8 h light/dark photoperiod, and a temperature of 19°C during the day and 15°C at night.

### Plant experiments under drought and well-watered

Twenty five plants were randomly divided into two distinct watering regimes: 10 plants for drought stress treatment and 15 for the well-watered control (“control” from here on). Sample collection took place at 14, 28, and 35 days post sowing, covering a pre-drought, a drought, and a post-drought time interval. The experimental timeline is presented in Figure S1. This experimental setup resulted in five biological replicates for each treatment and sampling time point. All plants were cultivated under well-watered conditions for 14 days, after which the drought group underwent a drought treatment for an additional 14 days by withholding water. During non-drought periods and for well-watered plants, distilled water was administered every other day to maintain a soil moisture content of 17% throughout the experiment. For plants subjected to drought stress, water was reintroduced to the PVC pots at the end of the drought period (28 days post sowing) to facilitate plant recovery.

### Sampling of rhizosphere and rhizoplane-associated microbial communities

Root sample collection and compartment processing were performed as described by (Guan *et al*., 2024) to obtain and rhizoplane (the root surface) associated microorganisms. Samples for DNA extraction were immediately flash frozen in liquid nitrogen and kept on dry ice until storing at −80°C. Samples were freeze-dried (CoolSafe 100-9 Pro freeze dryer, LaboGene, Lynge, Denmark). All freeze-dried samples were stored at −20°C until DNA extraction. Root and shoot length were recorded at sampling. Subsequently, roots and shoots were dried at 60°C for 3 days before determining the dry weight.

### DNA extraction

Genomic DNA was extracted from 500 mg of rhizoplane samples using the FastPrep-24TM 5G bead-beating system (MP Biomedicals, Irvine, CA, USA) set at 6.0 m/s for 40 seconds and the FastDNA^TM^ SPIN Kit for soil (MP Biomedicals), strictly following the manufacturer’s instructions. The DNA extracted from all samples was eluted in 60 μl elution buffer. The DNA’s purity and concentration were evaluated with a NanoDrop ND-1000 spectrophotometer (Thermo Fisher Scientific, Carlsbad, CA, USA). All DNA samples were stored at −20°C for subsequent quantitative PCR analysis and preparation of amplicon libraries.

### Chlorophyll quantification

The chlorophyll content of leaves of control and drought-stressed plants was determined at 28 days post sowing using a modified version of Liang’s method (Liang *et al*., 2017). The chlorophyll content was then calculated employing Arnon’s classic equations (Arnon, 1949) with absorbance values measured at 663 nm and 645 nm (full description in supplementary material).

### Determination of Superoxide Dismutase (SOD and Peroxidase (POX) and) activities in wheat leaves

SOD and POX activities were measured at 28 days post sowing, following a slightly modified version of the method previously described (Prochazkova *et al*., 2001). **SOD activity** was measured by recording the reduction in the optical density at 560 nm of the nitro-blue tetrazolium (NBT) dye catalyzed by the enzyme. **POX activity** was assessed based on the increase in optical density at 470 nm resulting from the formation of tetra-guaiacol. We refer to the supplementary material for full details.

### Strains and growth conditions

The viscosin-producing model strain *P. fluorescens* SBW25, along with its mutant strain that is deficient in viscosin production, was labelled using mCherry in a previous study (Guan *et al*., 2024). These strains were recovered from frozen glycerol stocks and cultured on Luria Broth Agar (LBA) plates, which consist of 1% tryptone, 0.5% yeast extract, 1% NaCl, and 1.5% agar, all supplemented with 10 μg ml^-1^ gentamicin. The plates were then incubated at 28°C for 48 h. Subsequently, single colonies were transferred into 20 ml liquid LB culture tubes with 10 μg ml^-1^ gentamicin supplementation and incubated at 28°C with continuous shaking at 180 rpm.

### Preparation of bacterial suspension and inoculation on wheat seedlings

The bacterial cells from the overnight culture were obtained by centrifuging at 6000×g for five minutes to yield cell pellets. These pellets were subsequently washed twice and resuspended in 0.9% sterile NaCl. OD_600_ for each inoculum was standardized to 1.0, equivalent to approximately 5 × 10^8^ CFU ml^-1^. The roots of three-day-old germinated seedlings were immersed in either *P. fluorescens* SBW25 mCherry suspension (WT) or the *P. fluorescens* SBW25Δ*viscA* mCherry suspension ( Δ*viscA*) for two hours. Subsequent to the inoculation, all plants were cultivated as detailed above. Briefly, sampling and DNA extraction was performed as describe above.

### Quantitative PCR analysis

The quantification of WT and Δ*viscA* was executed using qPCR as described in (Guan *et al*., 2024). The specific primers utilized are listed in Table 2. The absolute abundance of the target *mCherry* gene was subsequently calculated based on a standard curve, derived from a ten-fold serial dilution of DNA from SBW25::Tn7::mCherry cells. This standard curve exhibited a dynamic range spanning a ten-fold dilution series from 10^2^ to 10^8^ copies μl^-1^, with three technical replicates for each dilution. The efficiency ranged from 99.2 to 99.8%, and R^2^ values were > 0.99 for all standard curves. In addition, the qPCR setup included three technical replicates of a non-template (nuclease-free water) negative control.

**Table 2.**
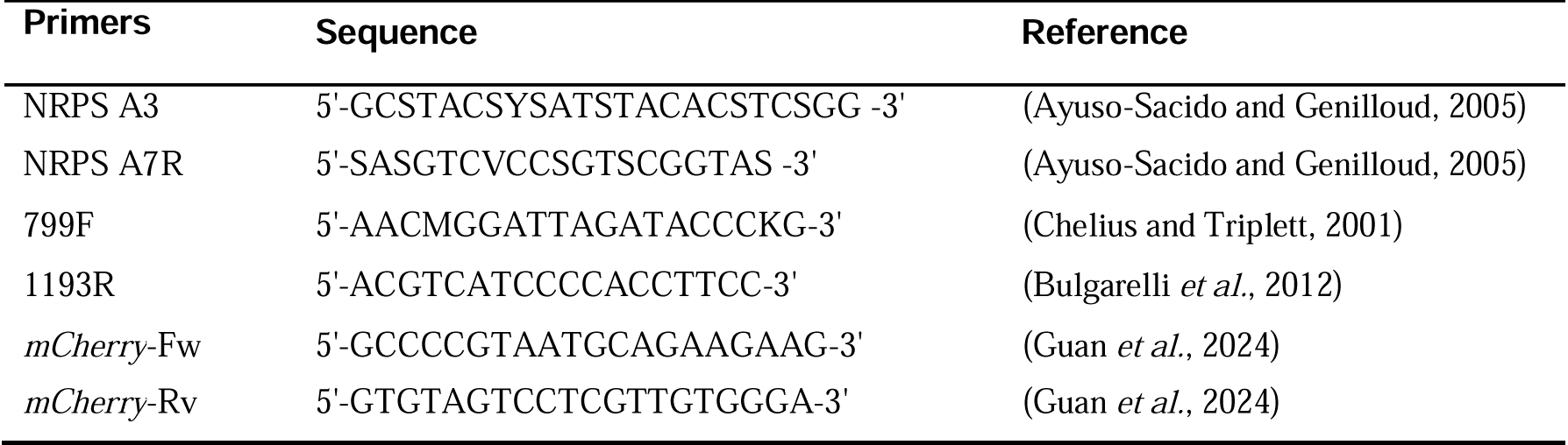
Primers used in the study.

### NRPS A domain and 16S rRNA amplicon sequencing

Primers A3F and A7R, detailed in Table 2, were used in a PCR reaction to amplify the NRPS A domains (Ayuso-Sacido and Genilloud, 2005). A two-step PCR and a dual-indexing approach were deployed for Illumina MiSeq sequencing. The PCR amplicons were produced using 15 µl of Platinum II Hot-start PCR Master Mix (Thermo Fisher Scientific), 0.6 µl of forward and reverse primers each (10 μM), 8.8 µl of nuclease-free water, and 5 µl of template. The PCR thermocycler protocol encompassed an initial denaturation phase at 95°C for 2 minutes, succeeded by 33 cycles of 95°C for 15 seconds, 55°C for 15 seconds, and 72°C for 15 seconds, with a final elongation step at 72°C for 5 minutes. The PCR amplification was verified via 1.5% agarose gels with loading buffer, and the PCR products were subsequently purified using AMPure XP beads (Beckman Coulter Inc. Brea, CA, USA). The purified DNA concentration was reassessed with a NanoDrop ND-1000 spectrophotometer (Thermo Fisher Scientific, Carlsbad, CA, USA) and a Qubit 2.0 fluorometer using the High-Sensitivity DNA assay (Thermo Fisher Scientific). Finally, the subsequent library construction and Illumina MiSeq sequencing (2×300 bp) were undertaken by the Erasmus Center for Biomics (Rotterdam, Netherlands).

The V5-V7 region of the bacterial 16S rRNA gene was amplified using primers 799F (Chelius and Triplett, 2001) and 1193R (Bulgarelli *et al*., 2012) (Table 2). Notably, this primer pair results in minimal amplification of plant mitochondria and chloroplast DNA (Beckers *et al*., 2016). The ZymoBIOMICS Microbial Community DNA Standard (Zymo Research, Irvine, CA, USA), pure culture SBW25 DNA, and water negative controls were included. A two-step dual-indexing approach was deployed for Illumina MiSeq sequencing, as described in (Guan *et al*., 2024). The subsequent library construction and Illumina MiSeq sequencing (2×300 bp) was performed by Eurofins Genomics (Ebersberg, Germany).

### Sequence processing

The raw amplicon sequence reads were processed using the DADA2 pipeline (version 1.30) (Callahan *et al*., 2016). Aside from a few parameter alterations, all other settings were retained as default. Specifically, for 16S rRNA gene reads, based on sequence quality, filtering was applied using default parameters, except for ‘trimLeft’ and ‘truncLen’. Primers were removed using ‘trimLeft’ = 19/18, depending on primer length, and reads were truncated using ‘truncLen’=280/200 for the forward and the reverse reads, respectively avoid poor quality and ambiguous sequences. Chimeras were removed after merging denoised pair-end sequences. Each unique 16S rRNA amplicon sequence variant (ASV) was classified according to the SILVA database (version 138.1) (Quast *et al*., 2013). Non-bacterial ASVs, including chloroplasts and mitochondrial reads, were filtered out, and the mock community and negative controls were checked for contamination. For NRPS A domains reads, solely the forward reads were employed, as recommended in (Tracanna *et al*., 2021). Reads were truncated at 250 nt, and quality filtered using maxEE = 7. NRPS A domain reads were annotated using a modified version of the dom2bgc pipeline (Tracanna *et al*., 2021). In brief, forward NRPS reads were translated into six frames and aligned the reads. Translated protein sequences were clustered based on Euclidean distances (Tracanna *et al*., 2021) and annotated using the most recent versions of MiBig (v. 3.1) (Terlouw *et al*., 2023) and antiSmash databases v.7.0.0 (Blin *et al*., 2023). In addition to the annotation performed using the dom2bgc pipeline, we performed a BlastP search of the NRPS ACs against the most recent MiBig database v.3.1 (Terlouw *et al*., 2023) with a threshold of the E value (10^-20^).

### Data analysis and statistics

Statistical analysis was performed in R version 4.3 (R Core Team, 2020). Differences in plant parameters and gene copy numbers between the drought stressed and control plants were analyzed by unpaired t-test (P < 0.05). P values were adjusted for multiple comparisons using “BH”. ANOVA was used to test for differences among different inoculations (P < 0.05).

For microbiome diversity and composition analyses, we used the R packages phyloseq version 1.46 (McMurdie and Holmes, 2013) and ampvis2 version 2.8.9 (Andersen *et al*., 2018). The amp_rarecurve function in the ampvis2 package was used to generate rarefaction curves for each sample. For determination of alpha diversity, richness, beta diversity and unique NRPS ACs, data was rarefied to an even depth using the mean values of 100 iterations. The alpha diversity and richness for the 16S rRNA and NRPS amplicons were determined using the Shannon diversity and CHAO1 indices with phyloseq_estimate_richness. Significance testing of the Shannon diversity was done with a t-test. For the NRPS A domain amplicons, the sample I23.1089.E04 was excluded from the analysis due to low read number.

A Principle Coordinates Analysis (PcoA) was performed using ordinate() in phyloseq (based on vegdist() from vegan) with rarefied data and Bray-Curtis dissimilarity matrix. Permutational Multivariate Analysis of Variance (PERMANOVA) based on Bray-Curtis dissimilarity matrix was used to test the effects of different factors (Sampling time, watering treatment) and their interactions on the beta diversity of communities in vegan version 2.6.6.1 (Oksanen *et al*., 2020). Dissimilarity matrices of 16S rRNA amplicons and NRPS amplicons were compared with mantel test (Pearson correlations) in vegan. The differential abundance of ASVs between groups was determined using beta-binomial regression with the corncob package version 0.4.1 (Martin *et al*., 2020). Only ASVs or ACs that had an estimated differential abundance of −1 or >1, and *P*-values adjusted for multiple testing < 0.05 (FDR < 0.05) were considered significant. We searched for ACs unique to either the drought stressed or control plants among the core ACs in the rhizoplane. Here, we defined core ACs to be present in at least four out five replicates. The core ACs of one treatment were then compared to all ACs from the other to determine unique core ACs. Identification of NRPS known to be produced by *Pseudomonas* was done using the complete list of compounds presented in (Zhou *et al*., 2024).

## Data availability

All raw sequencing data used in this study had been deposited in the NCBI Sequence Read Archive (SRA) database under accession codes <u>PRJNA987606</u> (16S rRNA) and <u>PRJNA987694</u> (AD). The scripts describing data treatment is available at https://github.com/Edmondbrn/Linking_microbial_community_structure_with_function.

## Acknowledgments

We acknowledge the assistance of Marnix H. Medema and Vittorio Tracanna in analyzing the NRPS A domain amplicon data. This study was funded by the Novo Nordisk Foundation (Grant number: NNF19SA0059360). YG expresses gratitude to the Chinese Scholarship Council for providing a PhD scholarship (CSC Grant 201908510124). FB was supported by the Independent Research Fund Denmark (10.46540/2031-00010B).

