## Supplementary material for "Linking microbial community structure with function using amplicon sequencing of NRPS genes associated with wheat roots under drought stress": Supp figures

**Supporting information for:**

Linking microbial community structure with function using functional amplicon sequencing of the NRPS genes around wheat roots during drought stress

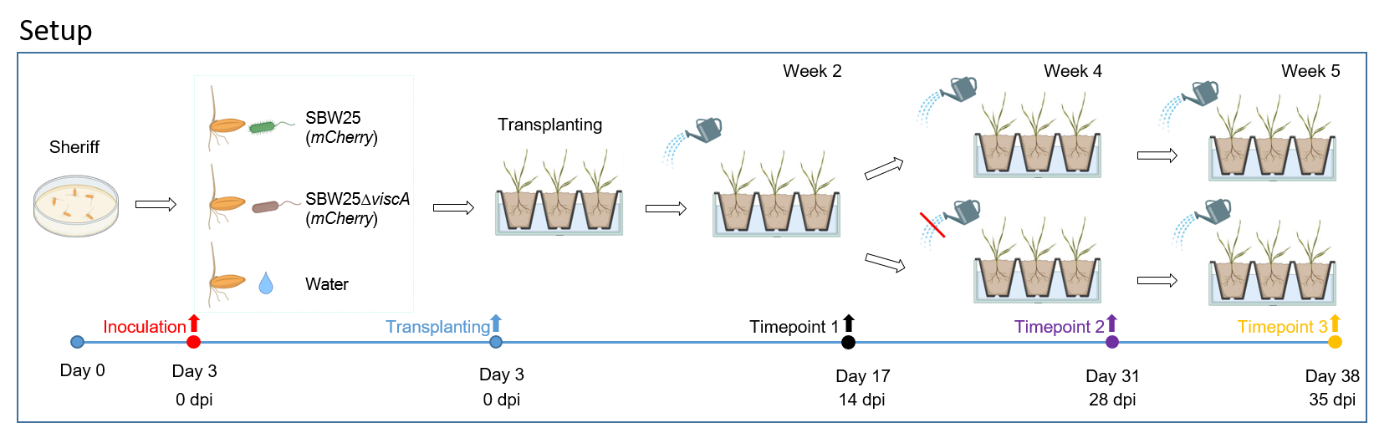

Figure S1. The timeline for plant experiments under drought and well- water conditions. Timeline is shown below, DPI (Days Post Inoculation).

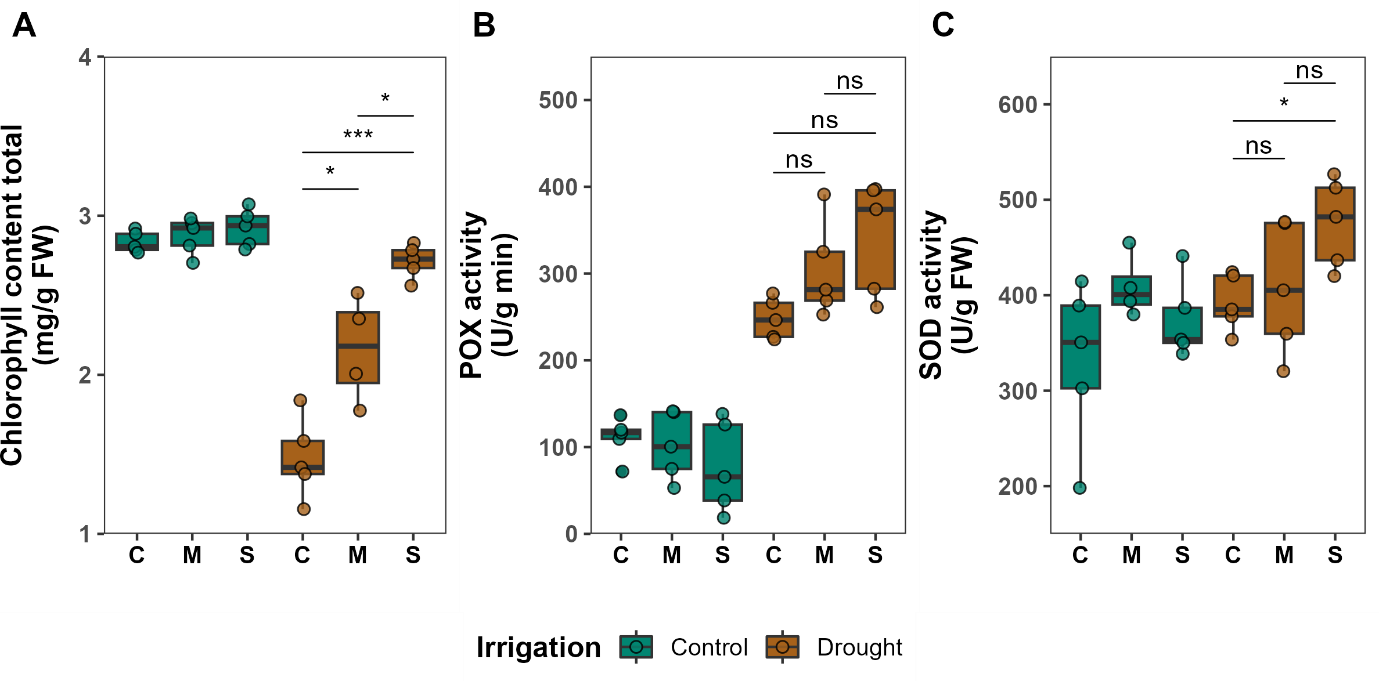
Figure S2. The effects of inoculation with SBW25 and mutant on chlorophyll content, POX and SOD activity were assessed 4 weeks post sowing drought stress and well-watered conditions. **A)** total chlorophyll content. **B)** POX activity. **C)** SOD activity. Each box plot represents data from five replicates. Each dot represents a sample point. The horizontal bars within boxes represent medians. The tops and bottoms of boxes represent the 75th and 25th percentiles, respectively. The upper and lower whiskers extend to data no more than 1.5× the interquartile range from the upper edge and lower edge of the box, respectively. C: Control (non-inoculated), M: ΔviscA Mutant, S: SBW25. ANOVA was used to determine whether inoculation was significant (p < 0.05) within irrigation. Asterisks indicate a statistically significant difference between the two inoculations (t-test) *p < 0.05 and **p < 0.01.

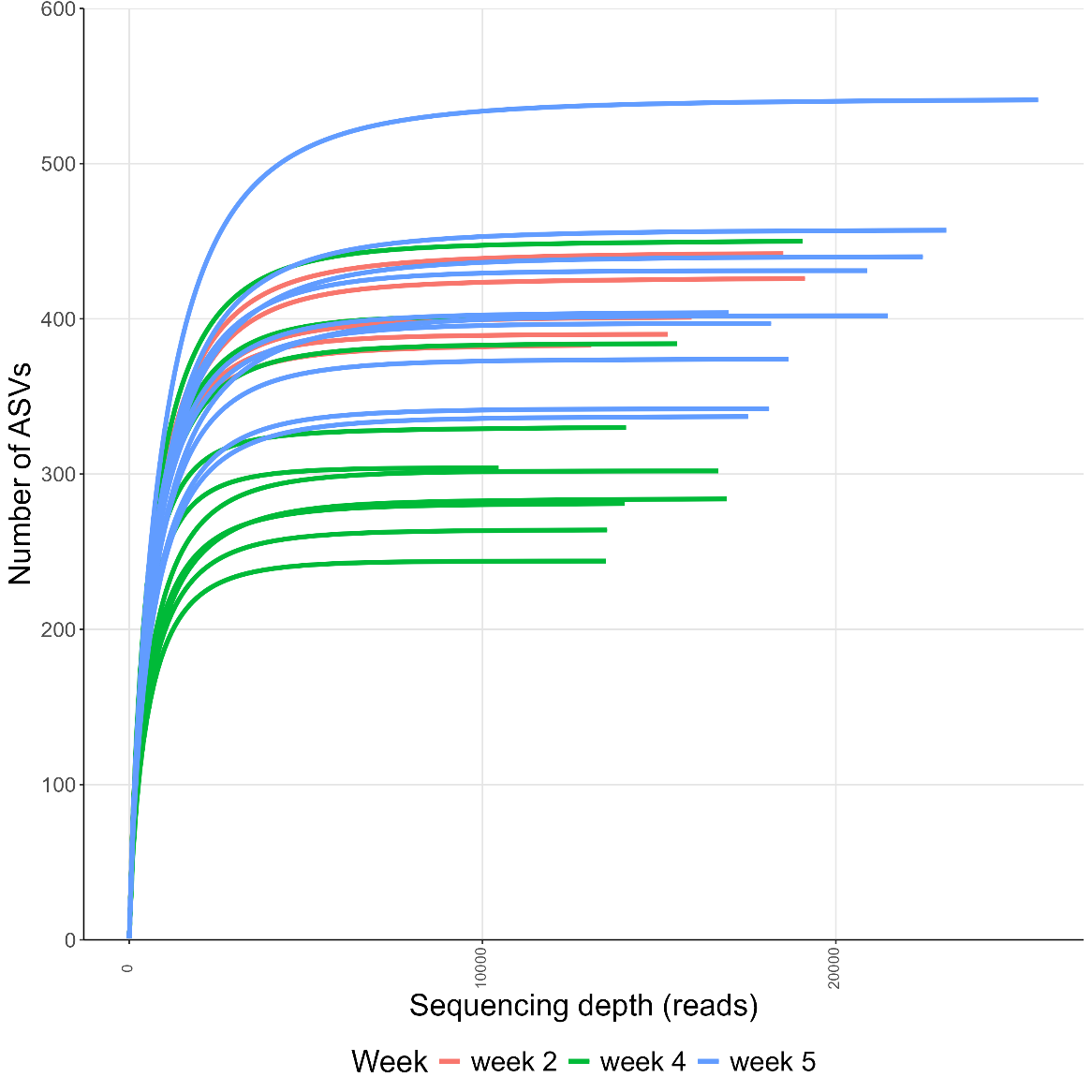

Figure S3. Rarefaction curves of the 16S rRNA samples.

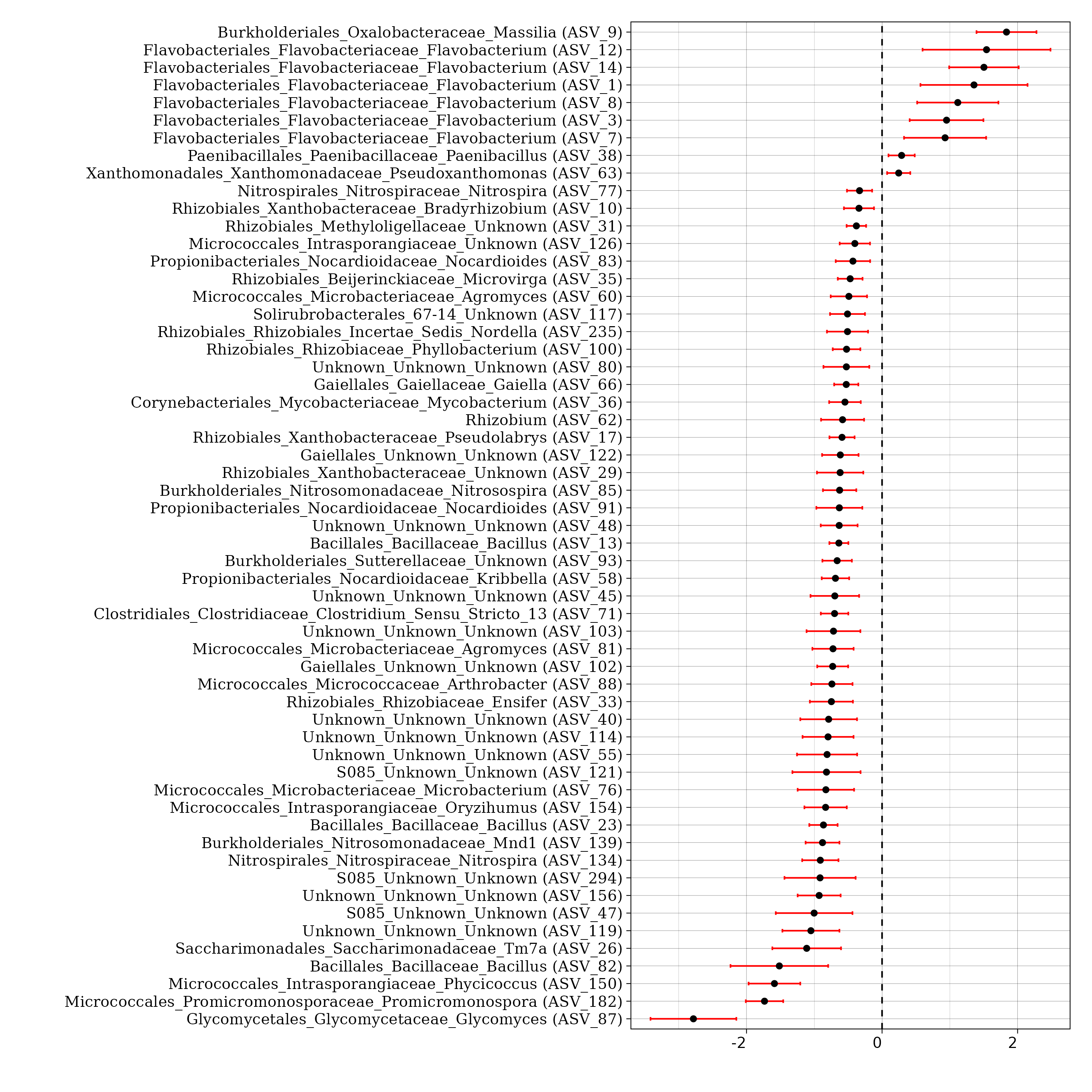

Figure S4. Differentially abundant bacterial ASVs between drought-stressed plants and control plants at week 4 determined by corncob, FDR < 0.05. Symbols on the right of the dashed line indicate an increase in relative abundance in the rhizoplane of the control plants compared to drought-stressed plants, whereas symbols on the left side indicate a higher relative abundance in drought-stressed plants.

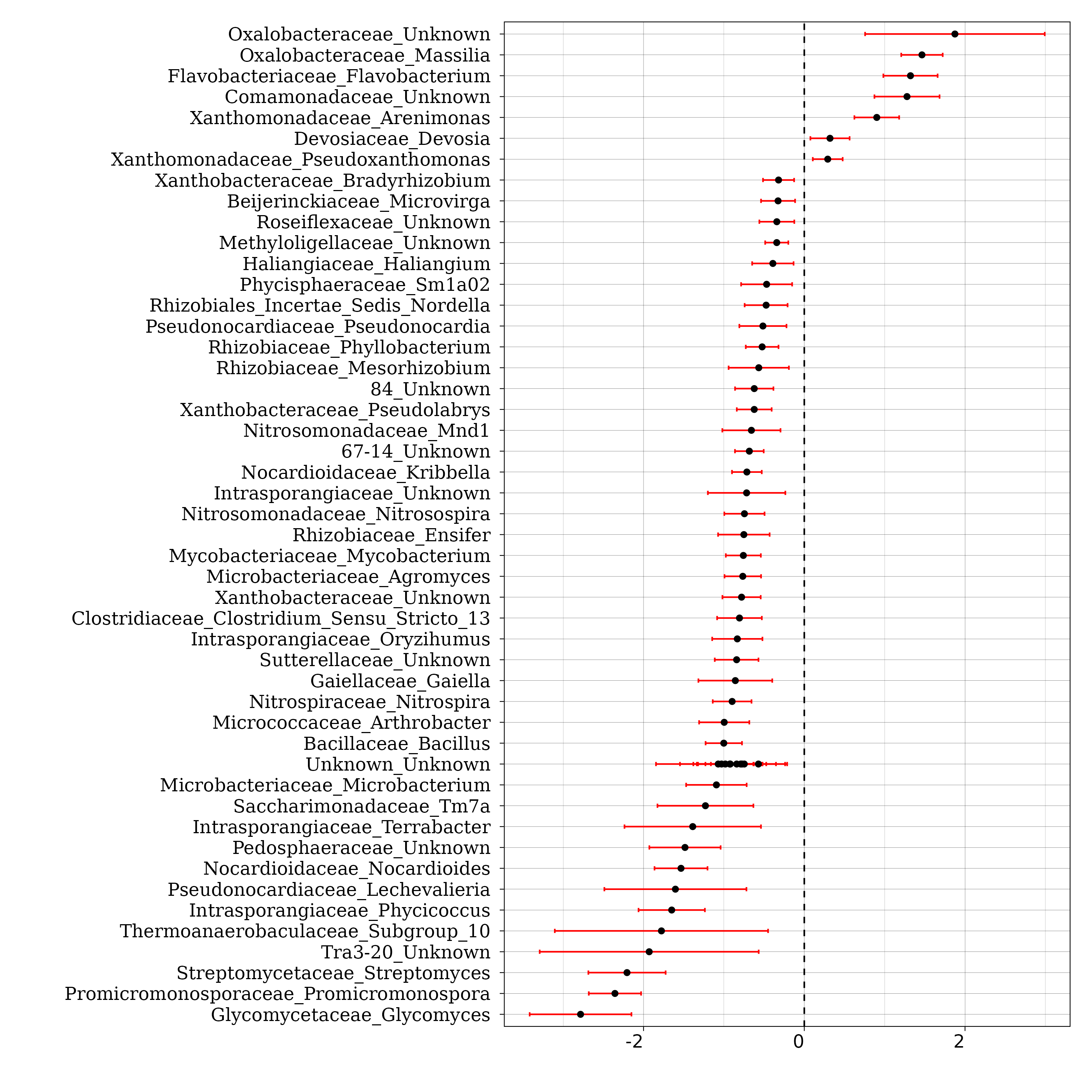

Figure S5. Differentially abundant genera between drought-stressed plants and control plants at week 4 determined by corncob, FDR < 0.05. Symbols on the right of the dashed line indicate an increase in relative abundance in the rhizoplane of the control plants compared to drought-stressed plants, whereas symbols on the left side indicate a higher relative abundance in drought-stressed plants.

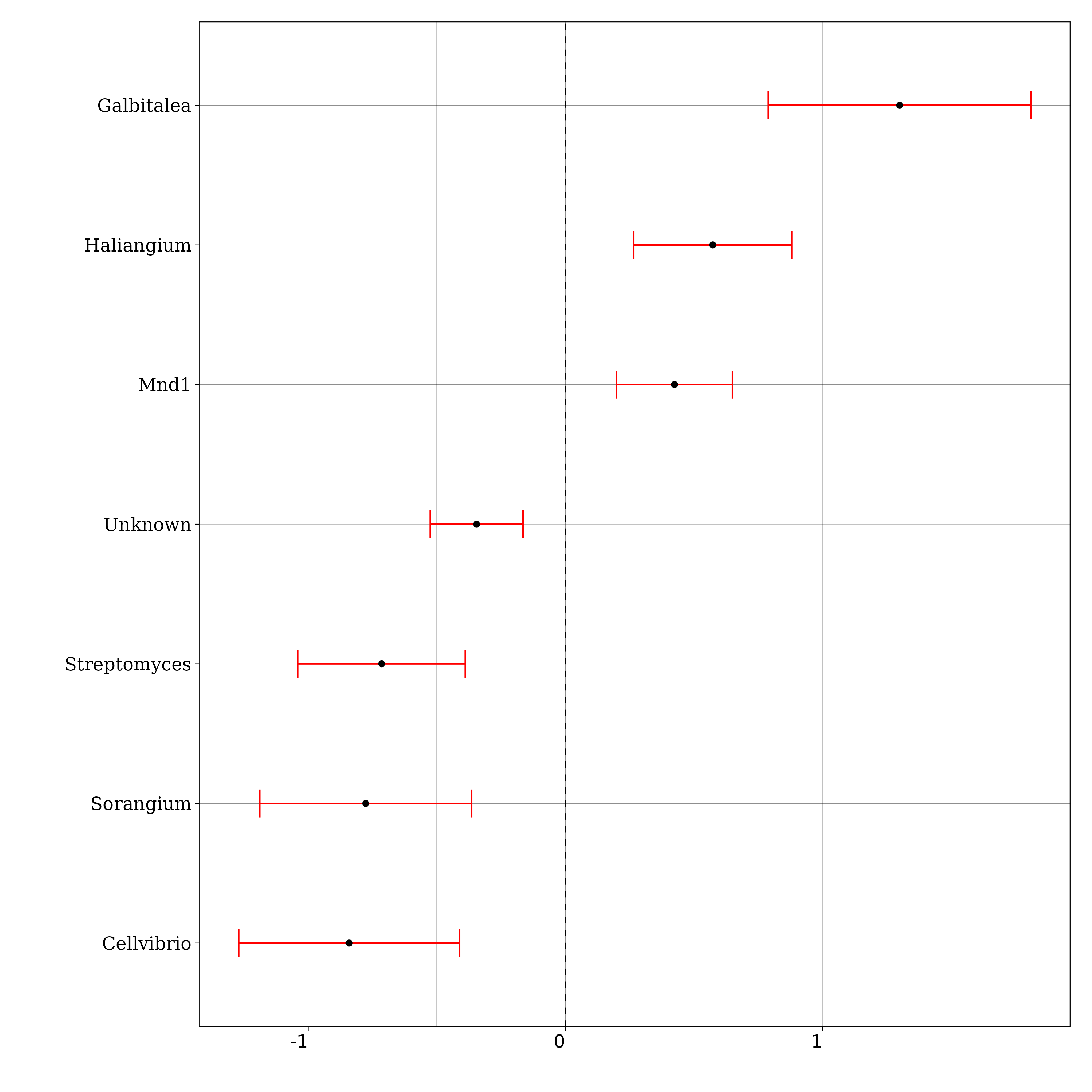

Figure S6. Differentially abundant genera between drought-stressed plants and control plants at week 5 determined by corncob, FDR < 0.05. Symbols on the right of the dashed line indicate an increase in relative abundance in the rhizoplane of the control plants compared to drought-stressed plants, whereas symbols on the left side indicate a higher relative abundance in drought-stressed plants.

| 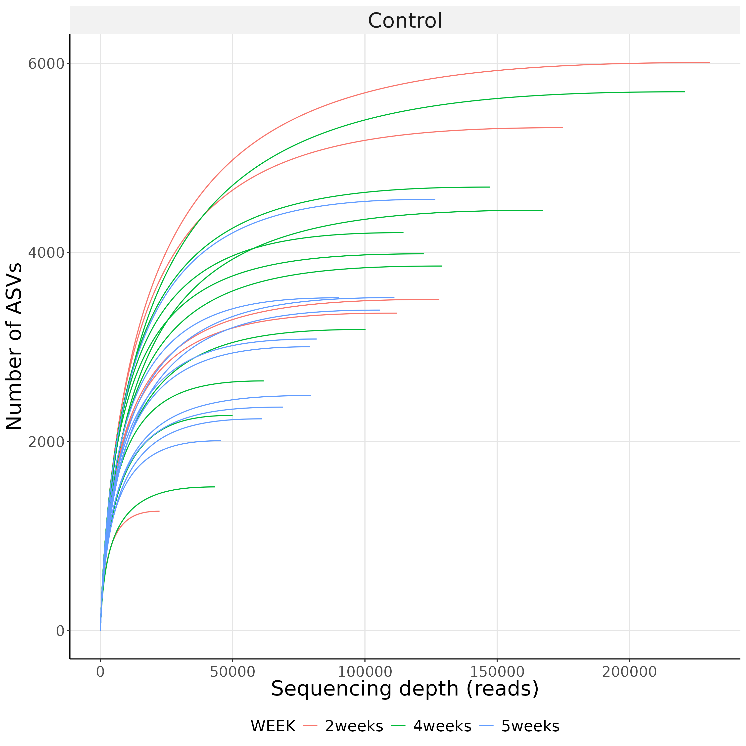 |
| --- |

Figure S7. Rarefaction curves of the NRPS samples rhizoplane.

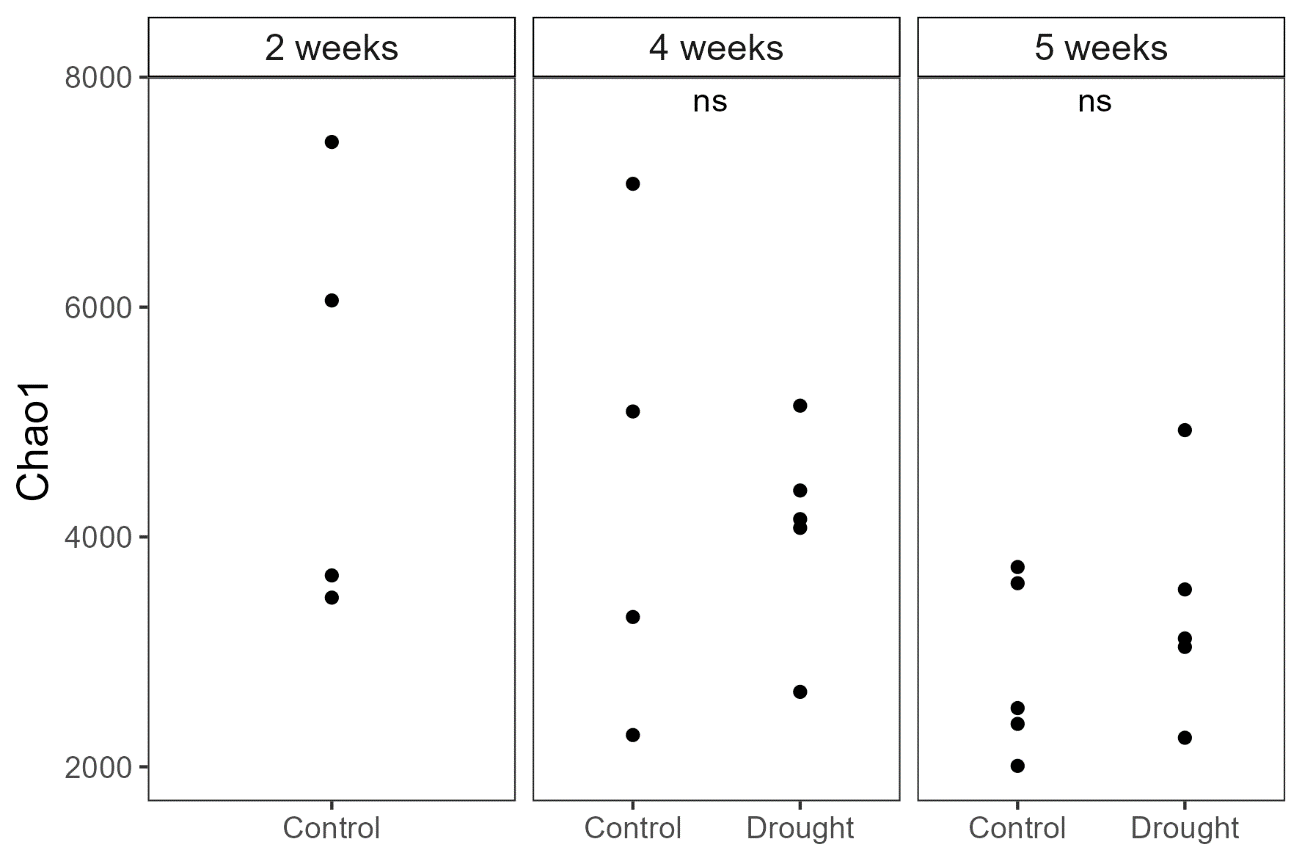

Figure S8. Chao1 richness of the NRPS ACs in the rhizoplane samples from control and drought stressed plants. NS = not significant (p > 0.05, t-test).

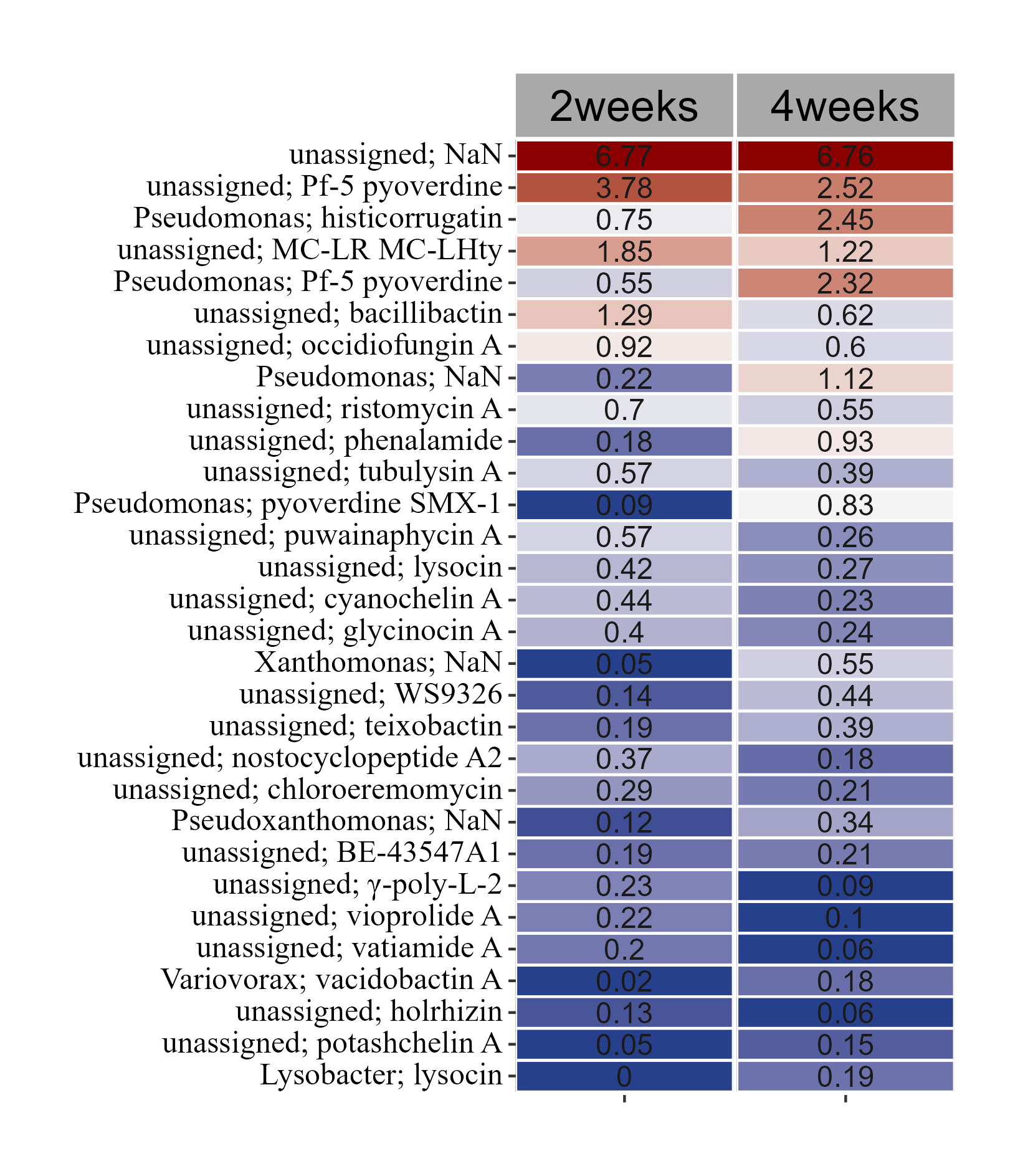

Figure S9. The 30 most abundant NRPS ACs which were differentially abundant between week 2 and 4. Differentially abundance was determined by corncob, FDR < 0.05. Values are mean relative abundances (n = 5).

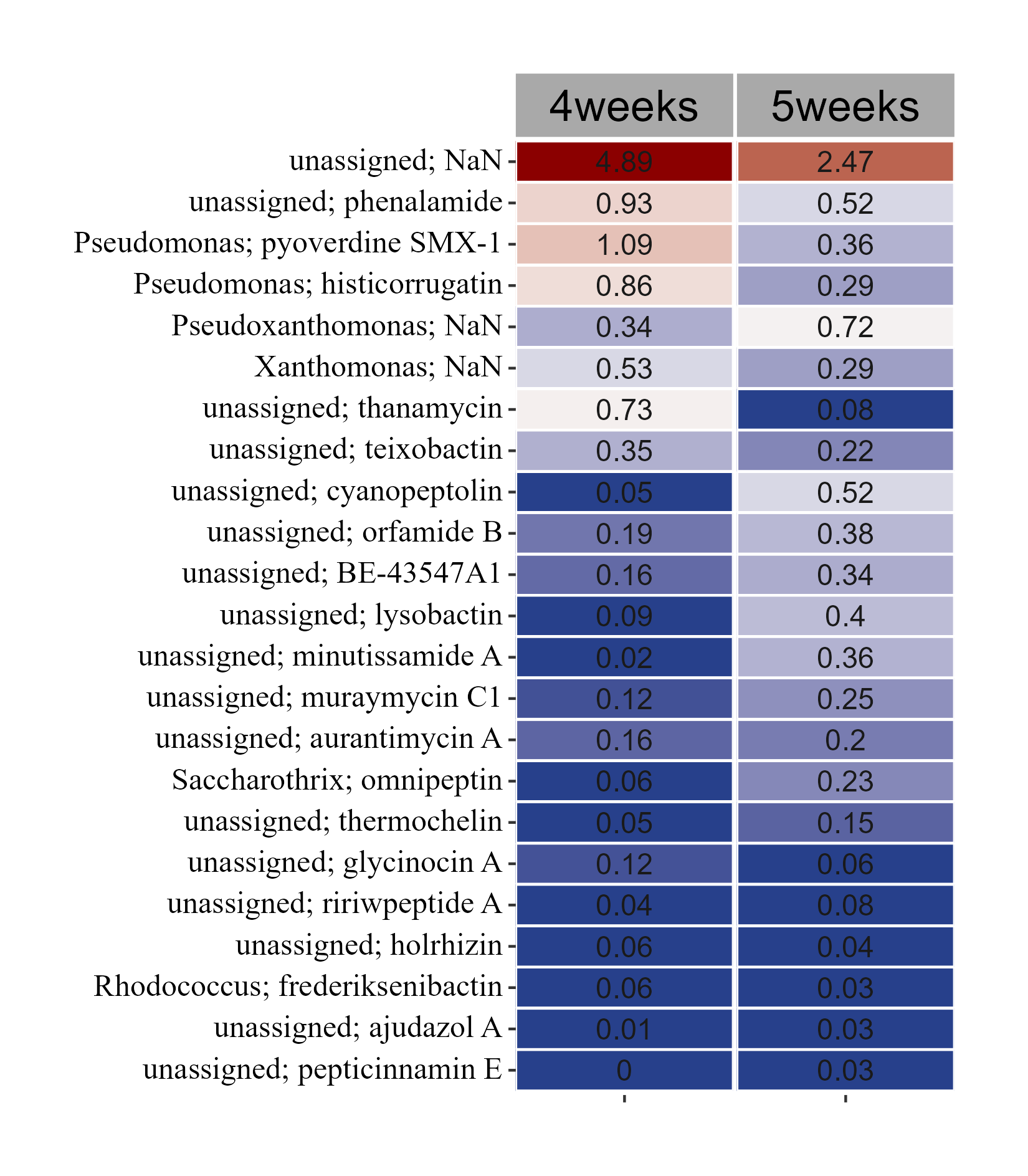

Figure S10. The 30 most abundant NRPS ACs which were differentially abundant between week 4 and 5. Differentially abundance was determined by corncob, FDR < 0.05. Values are mean relative abundances (n = 5).

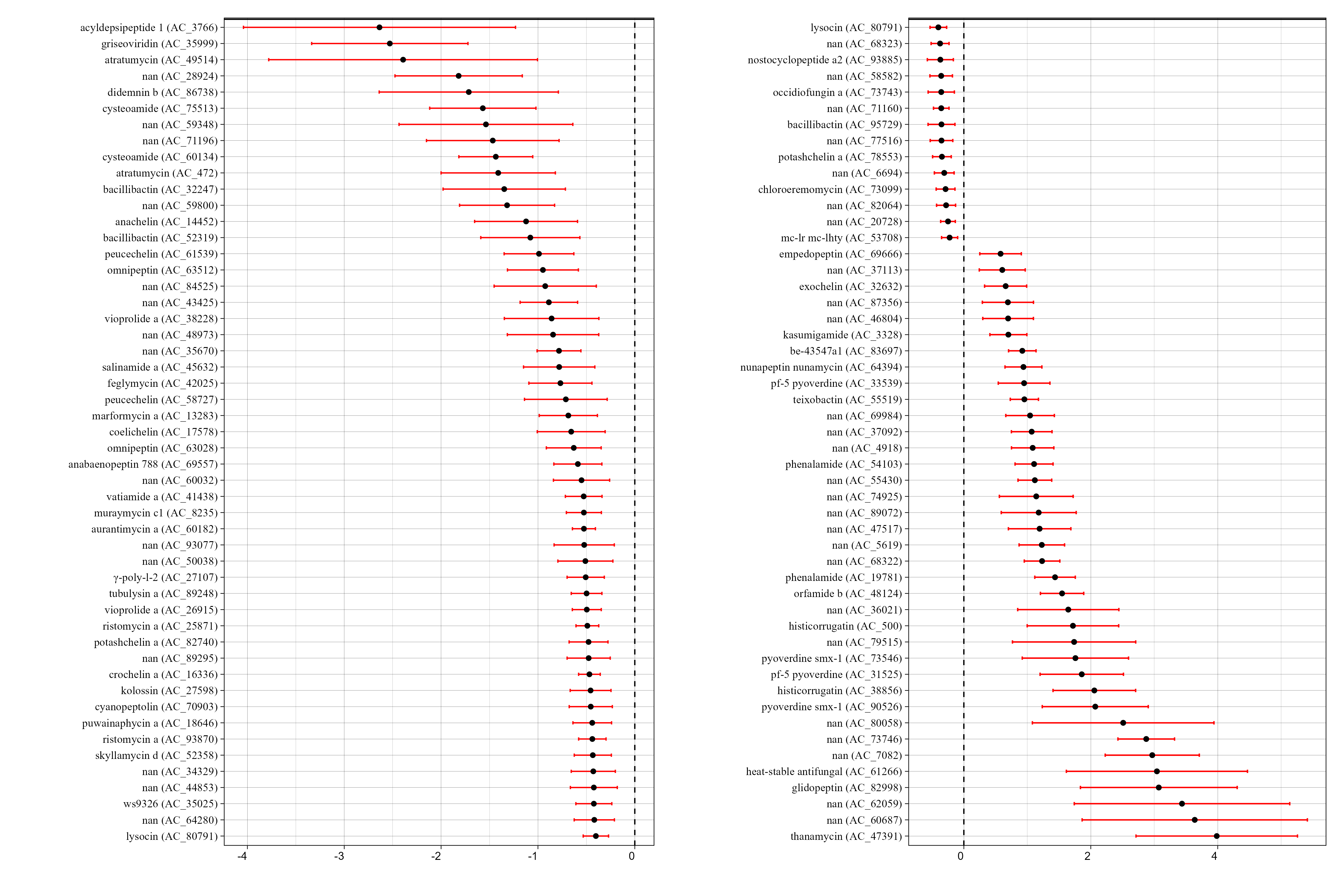

Figure S11. Differential abundant ACs between drought-stressed plants and control plants at week 4 determined by corncob, FDR < 0.05. Symbols on the right of the dashed line indicate an increase in relative abundance in the rhizoplane of the control plants compared to drought-stressed plants, whereas symbols on the left side indicate a higher relative abundance in drought-stressed plants.

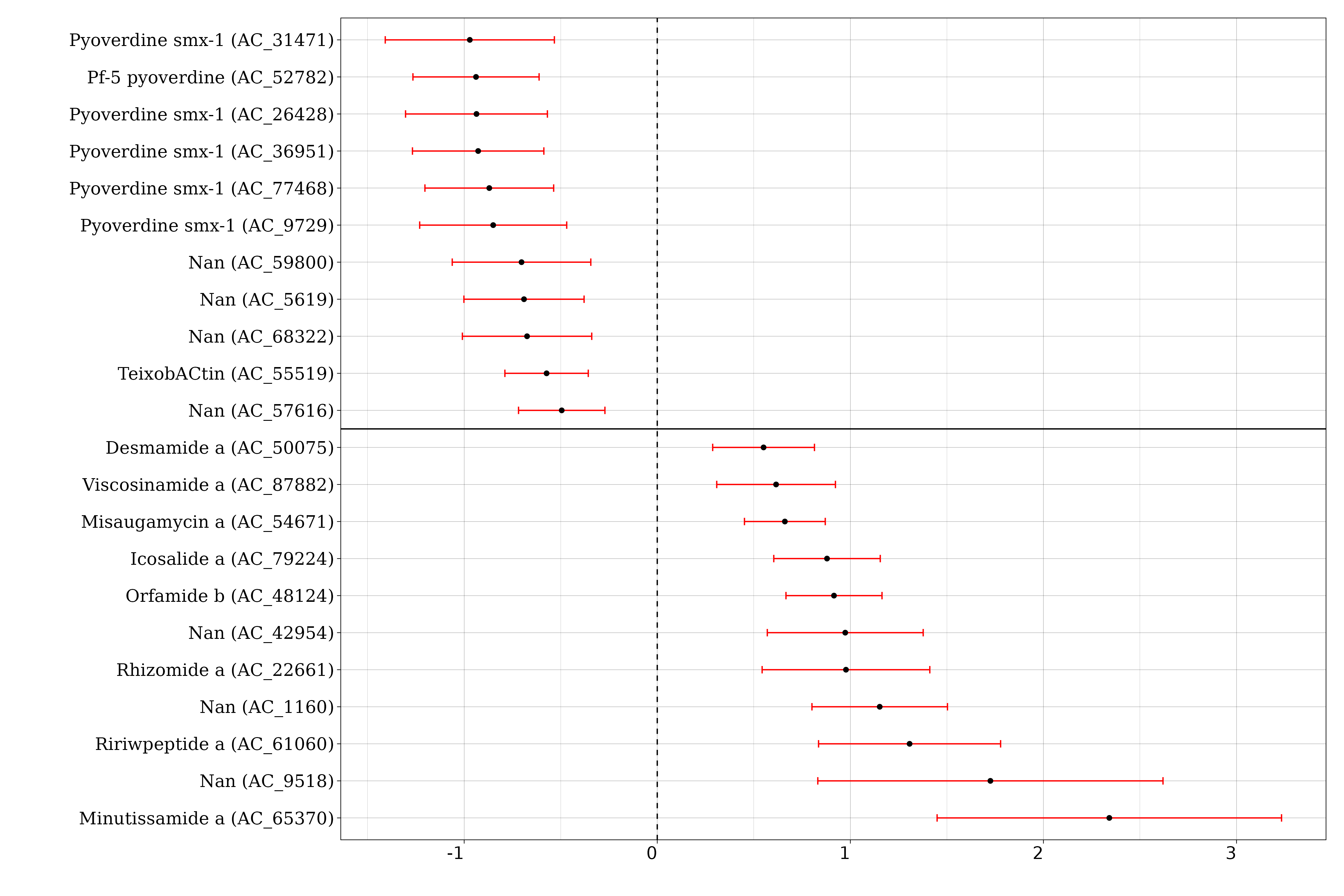
Figure S12. Differential abundant ACs between drought-stressed plants and control plants at week 5 determined by corncob, FDR < 0.05. Symbols on the right of the dashed line indicate an increase in relative abundance in the rhizoplane of the control plants compared to drought-stressed plants, whereas symbols on the left side indicate a higher relative abundance in drought-stressed plants.

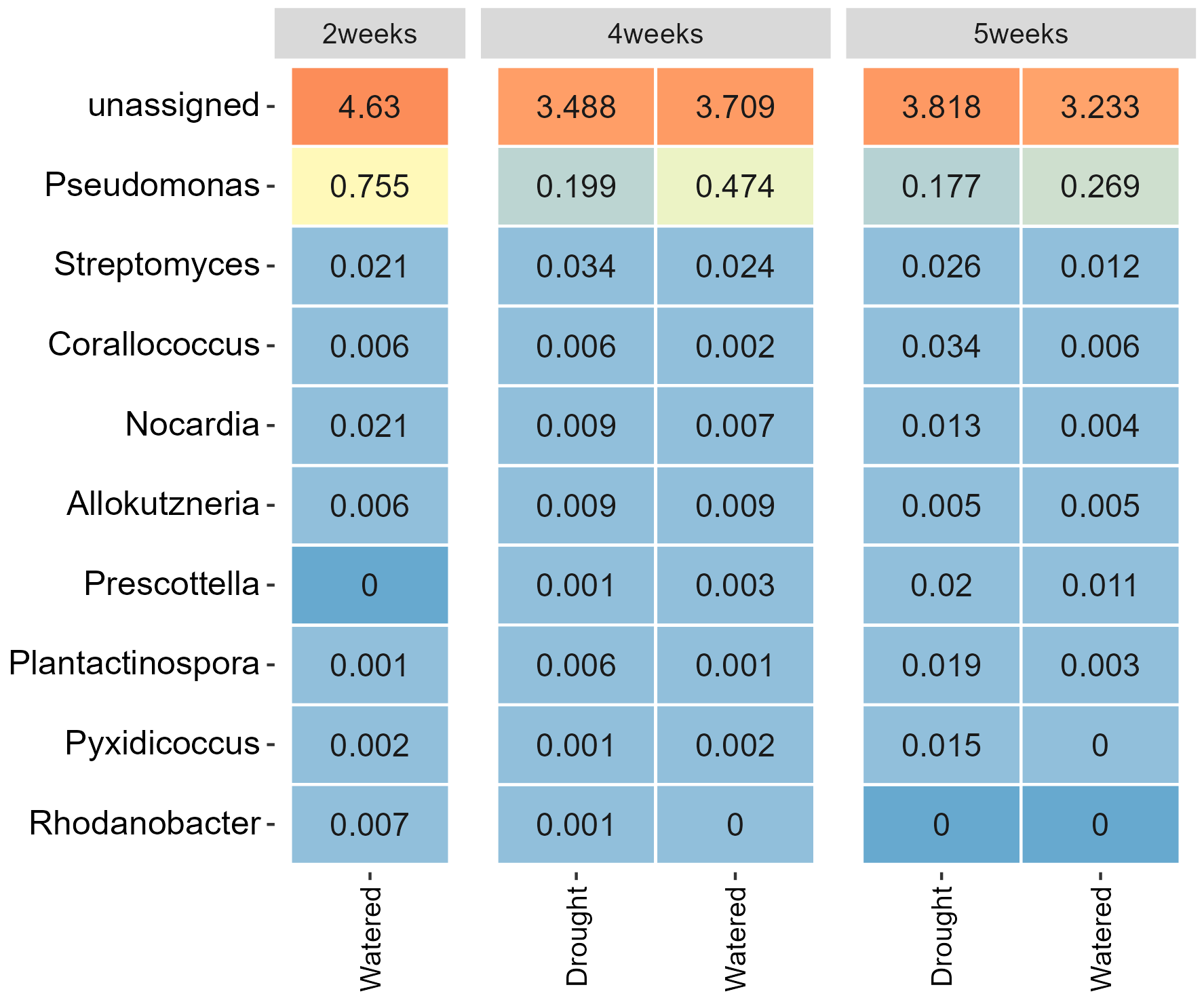

Figure S13. The relative abundances of the genera with the highest proportion of CLPs based on MiBig blast results. The relative abundance is of all NRPS ACs.
