## supporting methods for "Linking microbial community structure with function using amplicon sequencing of NRPS genes associated with wheat roots under drought stress"

**Chlorophyll quantification**

The chlorophyll content of leaves of both well-watered and drought-stressed plants was evaluated at 28 dpi using a modified version of Liang's method (Liang *et al.*, 2017). For the assessment, an exact weight of 0.1 g of leaf tissue was obtained, followed by finely chopping the tissue and immersing it in 10 ml of 80% acetone. These samples were then stored in darkness for 48 hours and subsequently centrifuged at a speed of 14,000 g for 5 minutes to achieve a clear mixture. The chlorophyll content was then calculated employing Arnon’s classic equations (Arnon, 1949) with absorbance values measured at 663 nm and 645 nm. The equations used are as follows:

Total chlorophyll (mg/L) = 20.21 (A645) + 8.02 (A663)

Chlorophyll content (mg/g) = C×V/A×1000

Where:

C: chlorophyll concentration in mg/L

V: total volume of extraction solution in ml

A: fresh weight of the sample in g

**Determination of Peroxidase (POD) and Superoxide Dismutase (SOD) activities in wheat leaves**

SOD and POD activities were measured at 28 dpi, following a slightly modified version of the method previously described (Prochazkova *et al.*, 2001). An exact weight of 0.1 g of leaf material was ground into a homogenous suspension with 2 ml of cooled extraction medium (phosphate buffer with 1% polyvinylpyrrolidone, pH 7.8) in a chilled mortar. The extraction medium was used to rinse the mortar, bringing the final volume to 10 ml. After centrifuging the mixture at 4°C, 10000 rpm for 15 minutes, the resulting supernatant was utilized as the enzyme source.

SOD activity was estimated by recording the reduction in the optical density of the nitro-blue tetrazolium (NBT) dye catalyzed by the enzyme. Three milliliters of the reaction mixture, comprising 130 mM methionine, 0.75 mM nitroblue tetrazolium chloride, 0.1 mM EDTA-Na_2_, 50 mM phosphate buffer (pH 7.8), and 0.1 ml enzyme, was initiated by adding 20 μM riboflavine and placing the tubes under fluorescent lamps for 25 minutes. The reaction was halted by switching off the light and storing the tubes in darkness. A complete reaction mixture without the enzyme, which yielded the maximum color, was used as a control, and a non-irradiated complete reaction mixture served as a blank. The optical density was documented at 560 nm, and a single unit of enzyme activity was considered as the quantity of enzyme that diminished the absorbance value by half relative to test tubes devoid of the enzyme.

POD activity was assessed based on the increase in optical density resulting from the formation of tetra-guaiacol. The 2.5 ml reaction mixture contained 3.4 mM guaiacol, 18 mM H_2_O_2_, 0.2 M phosphate buffer (pH 6.0), and 25 μl enzyme. The optical density resulting from the generation of tetra-guaiacol was measured at 470 nm, and a single enzymatic unit (u) was characterized as a 0.01 increment in the absorbance measurement per minute.
